# An umbrella review of randomized control trials on the effects of physical exercise on cognition

**DOI:** 10.1101/2022.02.15.480508

**Authors:** Luis F. Ciria, Rafael Román-Caballero, Miguel A. Vadillo, Darias Holgado, Antonio Luque-Casado, Pandelis Perakakis, Daniel Sanabria

## Abstract

Extensive research links regular physical exercise to an overall enhancement of cognitive function across the lifespan. Here, we assess the causal evidence supporting this relationship in the healthy population, using an umbrella review of meta-analyses limited to randomized controlled trials (RCTs). Despite most of the 24 reviewed meta-analyses reporting a positive overall effect, our assessment reveals evidence of low statistical power in the primary RCTs, selective inclusion of studies, publication bias, and large variation in combinations of preprocessing and analytic decisions. In addition, our meta-analysis of all the primary RCTs included in the revised meta-analyses shows small exercise-related benefits (*d* = 0.22, 95% CI [0.16, 0.28]) that became substantially smaller after accounting for key moderators (i.e., active control and baseline differences; *d* = 0.13, 95% CI [0.07, 0.20), and negligible after correcting for publication bias (*d* = 0.05, 95% CrI [−0.09, 0.14]). These findings suggest caution in claims and recommendations linking regular physical exercise to cognitive benefits in the healthy human population until more reliable causal evidence accumulates.

## Introduction

The physiological and health benefits of regular physical exercise are seemingly indisputable according to the scientific evidence accrued over the last century^1^. In addition, there has been a steady surge of studies reporting cognitive and brain benefits of regular physical exercise in healthy individuals across the lifespan^2^. These findings are driving current public health policies aimed at fostering exercise adherence^3^, in consonance with the World Health Organization, which currently recommends regular exercise as a means to maintain a healthy cognitive state^4^. One would therefore dare to say that the positive effect of chronic physical exercise at the cognitive level in the healthy population is nowadays taken for granted. The question we pose here is whether those claims, policies and recommendations are strongly supported by scientific evidence.

Summarized in many narrative^5^ and systematic reviews^6^ and a considerable number of meta-analyses^7–9^, the main conclusions of this literature are that: (1) the regular practice of physical exercise boosts cognitive performance in children, adolescents, and older adults, with limited evidence in young adults; (2) the impact seems especially prevalent in executive functions, although effects have also been described in other cognitive domains such as memory and attention; (3) the magnitude of the effects tends to be modest (*d* = 0.2–0.4), albeit reliable; (4) various factors might mediate and moderate the effects (e.g., exercise intensity, duration of the intervention, exercise training mode, etc.). This latter point is the principal caveat discussed in these articles, although the existence of the effect itself is rarely questioned.

Although they are not often highlighted, there are also reviews in the scientific literature reporting inconclusive evidence for a beneficial effect of physical exercise interventions on cognitive function in healthy populations. For example, in their meta-analysis Verburgh et al.^10^ found no evidence that physical exercise interventions had any effect in cognitively healthy older adults. The expert panel that recently conducted a systematic review on the topic^11^ also ended up stating that the evidence was inconclusive to claim that physical activity (a more general term that includes exercise) improves children’s cognitive or academic performance, except for the case of mathematics skills. Diamond and Ling^12^ claimed in a controversial article that the existing evidence shows that aerobic and resistance exercise training (arguably two of the exercise modes advocated as the best to improve cognitive performance^13^ are inefficient tools to enhance executive function.

The present umbrella review addresses the state of the art in this scientific topic by examining meta-analytic reviews limited to randomized controlled trials (RCTs), the current gold standard to ascertain causal links, to determine whether the claims regarding the benefits of regular physical exercise on cognition in healthy individuals are supported by solid and reliable empirical evidence.

## Results

Twenty-four meta-analyses^14,7,15,16,8,17–24,9,25–34^ meeting the inclusion criteria were selected among the 2,000 records retrieved in the search (see Fig. 1). We identified 271 primary studies in the meta-analyses, of which we selected 109 studies (Supplementary Table 1) that meet the inclusion criteria of our umbrella review (for details on exclusion criteria, see Supplementary Information). We extracted 737 effect sizes from the included primary studies (the information needed for effect size estimation was not available in seven primary studies^35–42^). The publication timeline of the reviewed meta-analyses and their respective primary studies reflects an exponential growth of the exercise–cognition topic in the last two decades (Fig. 2).

**Fig. 1.**
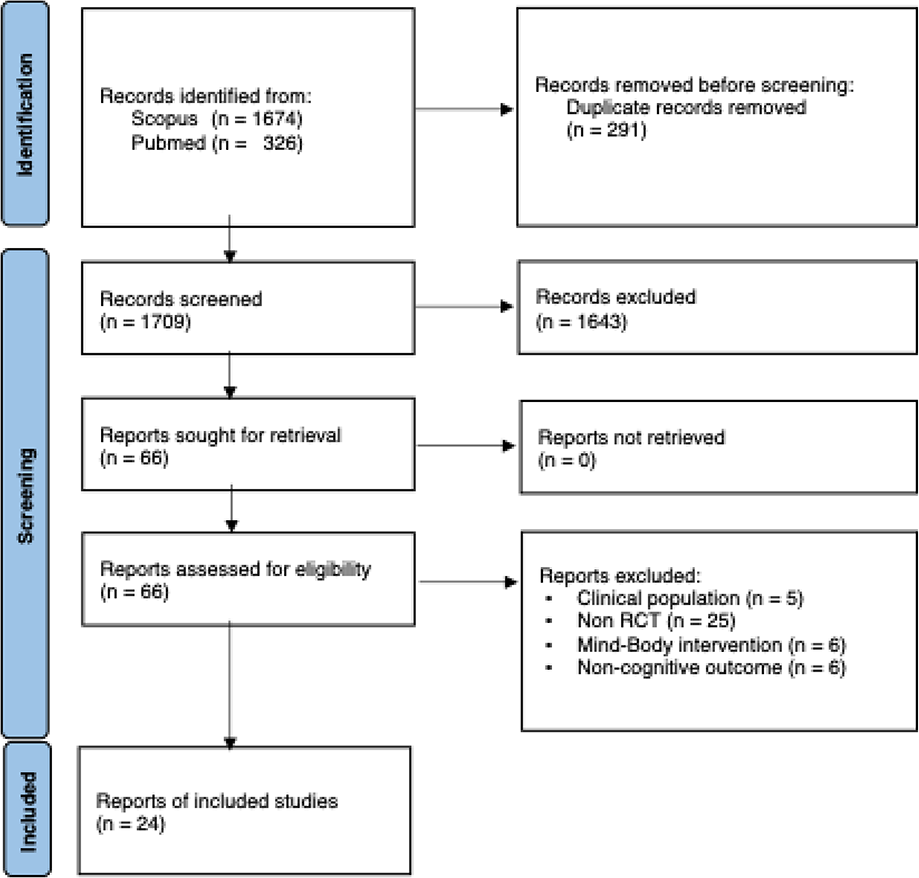
PRISMA flow chart for study inclusion. The initial search retrieved 2,000 records (Identification), among which 291 were removed as duplicates. After the screening of title and abstract, 66 records were chosen for full-article reading (Screening). Finally, 24 meta-analyses met the inclusion criteria of the umbrella review.

**Fig. 2.**
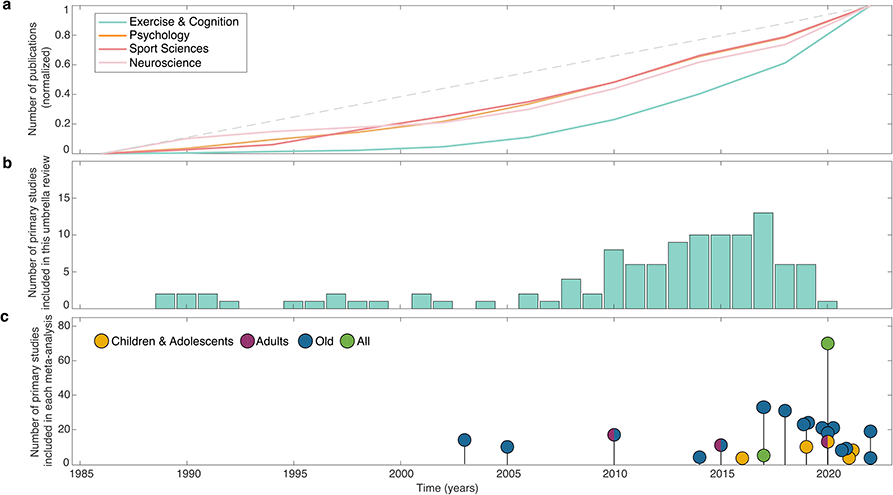
Evolution of the scientific literature. (A) Publication growth of Scopus-indexed articles in the areas of psychology (orange), sport sciences (red), neuroscience (pink), and exercise–cognition (turquoise) [using the search equation *(“sport” OR “exercise” OR “physical activity”) AND (“cognition” OR “executive functions” OR “executive control”)* in July 2022]. The number of publications per year was normalized to express the values of all the categories in a common scale from 0 to 1. Whereas the growth of the general categories closely followed a linear trend (gray dashed line), publications on the exercise– cognition topic depict an exponential proliferation highlighting the great interest generated by the topic in the last two decades. (B) Primary studies (RCTs) included in the present umbrella review. The 109 RCTs included show a similar exponential growth over the last 25 years, with a peak in 2017. (C) Timeline of meta-analyses included in this umbrella review along with the number of primary studies used by each meta-analysis to estimate the effect of regular exercise on cognition in healthy population. Dot color depicts the target age range of each meta-analysis.

Almost all of the 24 meta-analyses found a significant positive effect of regular exercise on cognition in healthy participants (median Cohen’s *d* = 0.29, 95% CI [0.23, 0.36]), concluding that exercise may improve cognitive skills (22^12,7,13,14,15–22,9,24–32^ out of 24). To avoid the influence of using non-target exercise programs or samples with health conditions, we analyzed the outcomes of primary studies only with target interventions and healthy populations. The re-estimated effects remained positive although disperse (median *d* = 0.17, 95% CI [0.13, 0.19]; range 0.06–0.39; Fig. 3A) and with variable heterogeneity (median *I*^2^ = 43.40%; range 0–96.91%; Fig. 3B). Part of the dispersion between the effects and heterogeneity could be due to the inclusion criteria adopted in each meta-analysis (Supplementary Table 1), differing in the type of physical exercise, age range of the participants, or cognitive outcome. Other sources of variability could come from the way the individual effect sizes were estimated or the strategy adopted in the meta-analyses to deal with the dependence generated by the inclusion of several outcomes from the same sample of participants. Further, sampling of primary studies and divergence in the inclusion criteria (see Variations in Study Sampling) could affect the results of the meta-analyses (the full assessment of the quality evaluation of each meta-analytic review included is available at the following link: https://osf.io/e9zqf/). Indeed, the number of primary studies included in each meta-analysis (excluding Lindheimer et al.^17^ with only one included study) ranged from 2 to 63 studies per meta-analysis (median of 11 studies; Fig. 3C). The median of the average number of participants per meta-analysis was 78 (range 36–673; Fig. 3D).

**Fig. 3.**
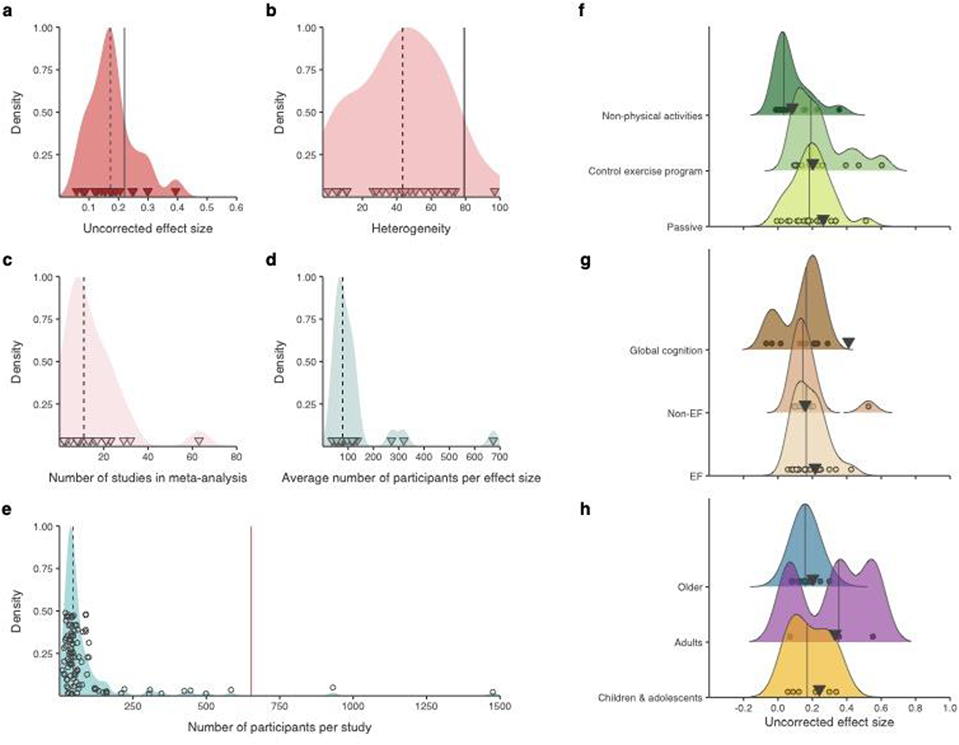
Reanalysis of the meta-analyses included in the umbrella review and influential variables. Distribution of (A) meta-analytic effect sizes re-estimated from the 24 meta-analyses and (B) their heterogeneity, expressed in *I*^2^. (C) Distribution of the number of studies, (D) the average number of participants per effect size in each meta-analysis, and (E) the total number of participants per primary study. Dashed lines indicate the average value for all the meta-analyses/primary studies, whereas black solid lines in (A) and (B) indicate the final effect and their heterogeneity in the full model (with the entire sample of primary studies). The red line indicates the required sample size for an acceptable power (see Methods for details). There was a small median effect size (*d* = 0.17, 95% CI [0.13, 0.19]) and substantial median heterogeneity (*I*^2^ = 43.40%). All except two primary studies used underpowered designs to assess a small effect size individually. Note that we excluded primary studies (from reviewed meta-analyses) involving mind-body, yoga (or similar), exercise programs combined with any other intervention (e.g., cognitive training), or samples of participants with medical conditions to avoid confounding factors. Therefore, the effect sizes represented might differ slightly from those originally reported by each meta-analysis. Estimated final effects of the included meta-analyses as a function of (F) the type of control program, (G) the type of cognitive outcome, and (H) the age range of the included samples. Vertical solid lines indicate the median of each category and the circles at the bottom the re-estimated effect of each meta-analysis, whereas the inverted triangles represent the full-model effect. Only the difference between passive controls and active controls was significant in the full model. However, this difference was not replicated with the medians of the re-estimated outcomes. Also, the median effect of young adults was larger than the other age cohorts showing a difference that did not appear when it was assessed in the entire sample of primary studies.

Since selective sampling of available evidence can influence the outcome of a meta-analysis, we subsequently assessed the effect of physical exercise on the entire sample of studies. We implemented a multilevel meta-analysis with all the primary studies included in the 24 meta-analyses. This full model served as reference estimate to interpret the findings in the reviewed meta-analyses. We observed an overall effect of *d* = 0.22, 95% CI [0.16, 0.28], *p* < .0001, and large heterogeneity, *I*^2^ = 79.52%. On average, the 24 originally reported effects and the 24 re-estimated departed 0.15 and 0.08 respectively from the summary effect of the full model. As might be expected, the number of included primary studies increased as a function of the year of publication (Kendall’s τ = 0.47, *p* < .001), in parallel to the exponential growth of primary articles. The number of participants per effect, in contrast, did not increase over the years (Kendall’s τ = 0.01, *p* = .913) and, more importantly, the number of participants were below the sample size required to achieve an acceptable statistical power (i.e., 80%) for a *d* = 0.22, the overall effect with the entire sample of primary studies (i.e., at least 652 participants; contrasting with a median of 48 participants, range 10–1476; Fig. 3D. See Methods for details on sample size estimation). At the primary study level, only two studies^43,44^ reached power equal to or higher than 80% (i.e., ≥ 652 participants; Fig. 3E).

### Variations in study sampling

The network visualization (Fig. 4A) of the reviewed meta-analyses reveals that the variability across meta-analyses might be in part due to divergences in the inclusion of primary studies. Altogether, the 24 meta-analyses comprise a total of 109 different primary studies, of which 28 (25.6%) were only included in one meta-analysis, and 21 (19.2%) in two of them. The scarce overlap is noticeable even between meta-analyses addressing the same age range and cognitive outcome. For instance, three meta-analyses focused on aging and general cognitive domains^22,23,25^ share only one primary study^45^ of the 23, 34, and 25 primary studies used, respectively. Although these three meta-analyses address the same topic with a similar approach (meta-analysis of RCTs) in a comparable time frame (between 2019 and 2020), their results are based on radically different bodies of evidence. The most recent meta-analysis included in this review^33^ deserves special attention. Although it addresses the same age target and cognitive outcome as the 3 meta-analyses mentioned above^22,23,25^, it does not share any primary studies with them. To further test this disconnection between meta-analytic reviews, we assessed the number of primary studies that were available at the time the meta-analyses carried out their last search and that met the inclusion criteria for each review (Fig. 4B). Overall, the meta-analyses included less than half of the available primary studies meeting their criteria (median of 48.77% of studies). Only six meta-analyses included most of the available studies (Aghjayan et al.^34^: 77%; Amatriain-Fernández et al.^31^: 80%; Angevaren et al.^7^: 83%; Colcombe & Kramer^14^: 73%; Haverkamp et al.^28^: 81%; Smith et al.^15^: 79%), whereas in a third of the meta-analyses, the analyzed studies represented less than 35% of all the potential targets^17,20,23,27,29,30,32,33^. This lack of overlap in the meta-analytic literature on the topic suggests that the conclusions drawn from these quantitative reviews cannot be taken as the empirical evidence accumulated over years, but as selective slices of it.

**Fig. 4.**
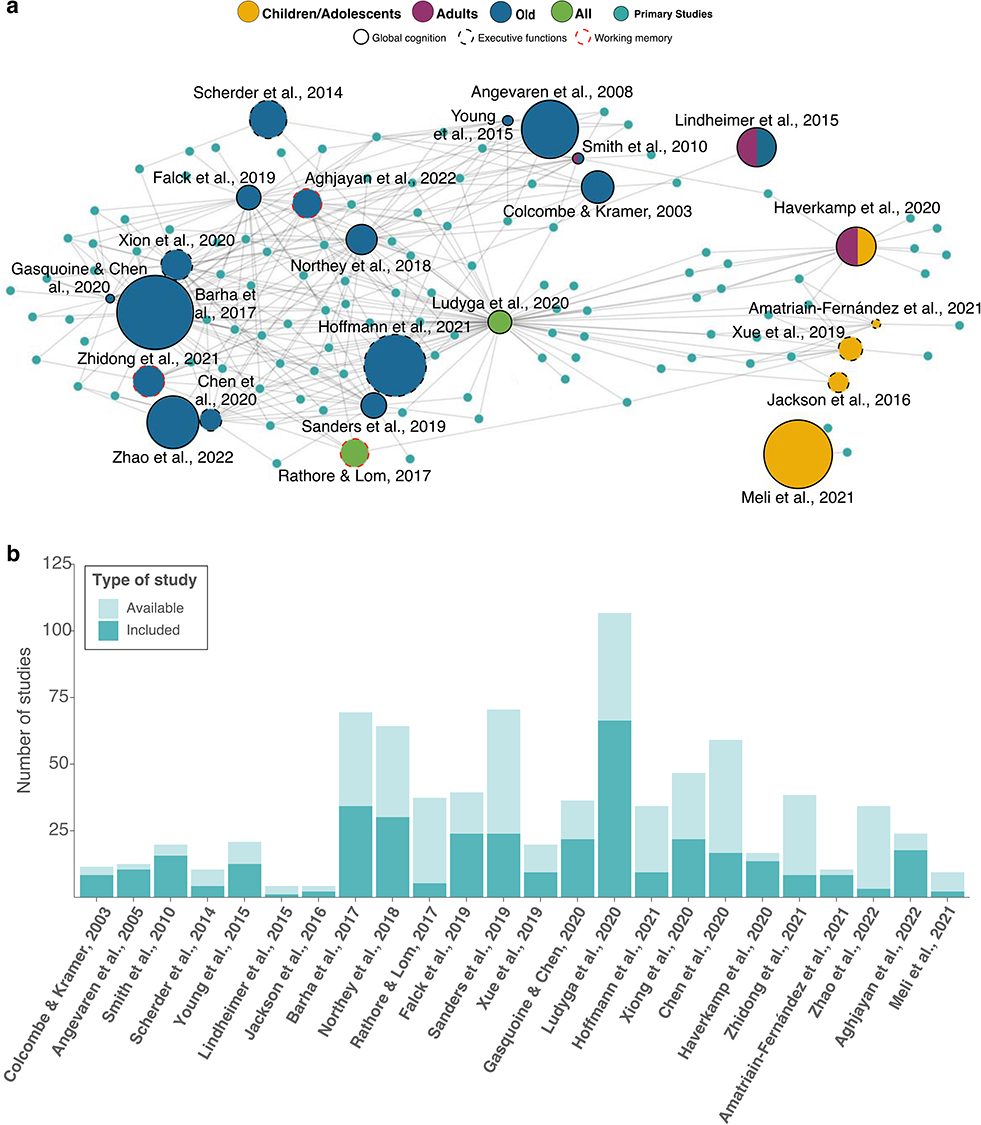
Network interaction among the meta-analyses included in the umbrella review. (A) Network of meta-analyses targeting RCTs on the relationship between regular physical exercise and cognitive functions in healthy populations included in this review. Meta-analyses are represented as nodes, with color indicating age range and edges depicting the cognitive domain addressed. The size of the nodes (for meta-analyses only) is proportional to the reported main effect size and the color indicates the age range addressed. Note that most meta-analyses report the effect of exercise on general cognitive domains, but some only report the effect of exercise on executive functions or memory. The primary studies used by each meta-analysis to estimate their reported effect size are represented by turquoise nodes (the size of these nodes is fixed). The inclusion of a primary study in a meta-analysis is represented by a gray line between the meta-analysis node and the primary study node. Spatially close meta-analyses share a greater number of primary studies, while more distant meta-analyses share few or none. It is important to highlight that this network visualization is basically descriptive and does not depict a complete picture of the field. It is intended to help clarify the state-of-the-art of the field rather than refute any particular hypothesis. (B) Number of primary studies included in each meta-analysis (darker turquoise bars) compared to the studies that met the specific inclusion criteria and were available at the moment when the last search of each meta-analysis was carried out (light turquoise bars). Meta-analyses are sorted by the date on which they conducted the last search for primary studies.

### Influential variables and publication bias

To check the influence of key moderating variables (i.e., type of control activity, type of cognitive outcome, age range, duration of the training programs, and baseline performance) we implemented separate meta-analytic models for each level of these variables using the entire sample of primary studies (i.e., full model). Whereas the effect was similar for all age cohorts (older: *d* = 0.20, 95% CI [0.12, 0.28]; adults: *d* = 0.33, 95% CI [0.01, 0.66]; children and adolescents: *d* = 0.24, 95% CI [0.12, 0.36]; all comparisons *p*s > .05) and all types of cognitive outcome (executive functions, EF: *d* = 0.22, 95% CI [0.15, 0.28]; non-EF: *d* = 0.16, 95% CI [0.09, 0.23]; global cognition: *d* = 0.41, 95% CI [0.00, 0.82]; all comparisons *p*s > .05), the observed benefit was larger with passive controls than with active control activities (*d* = 0.26, 95% CI [0.17, 0.35]; vs. non-physical active controls, *d* = 0.08, 95% CI [−0.01, 0.17], *p* = .007; vs. physical active controls, *d* = 0.20, 95% CI [0.08, 0.33], *p* = .426; Supplementary Figure 1a). The lack of statistically significant difference between age cohorts was observed even when the duration of the interventions was on average longer for older participants (median of 20 weeks, range 4–96) than with children and adolescents (median of 11 weeks, 6–96), and especially with young adults (median of 6 weeks, 4–24). Finally, the difference in performance between the groups at baseline also accounted for part of the variability. The benefits of physical exercise were greater when the performance at baseline in the experimental group was lower than the control group (β = −0.77, *p* = .011; Supplementary Figure 1b). Therefore, the type of control group and baseline differences could explain part of the between-study variability, reducing heterogeneity to *I*^2^ = 57.21%. The model including both moderators (i.e., control activity and baseline performance) predicted a decrease in the final effect to *d* = 0.13, 95% CI [0.07, 0.20], *p* < .001, in studies with active control and matched groups in baseline cognitive performance (i.e., pretest difference of *d* = 0). These findings with the full model were not replicated when we re-analyzed these variables in each meta-analysis within its set of primary studies. In the re-analysis we observed again substantial dispersion in the effects and, in some cases, the average estimates among meta-analyses do not correspond with the estimates of the full model (Fig. 3F–H).

Regarding publication bias, only 14^9,16,17,21–24,26–28,30,31,33,34^ out of 24 meta-analyses reported publication-bias analyses, most of them using funnel plot-based methods (i.e., visual inspection of the funnel plot, trim-and-fill procedure, and Egger’s method to test asymmetry in the funnel plot). Among those that assessed it, 7 found evidence of publication bias (7^9,22–24,26,28,34^ out of 14) and, among them, only 3 adjusted the final effect (3^9,22,34^ out of 7). Notably, one of the meta-analyses concluded benefits in episodic memory with physical exercise even after the publication-bias correction methods indicated a non-significant effect^32^.

We also tested the presence of publication bias and corrected the final effect in the meta-analyses with ten or more primary studies (12^8,9,15,19,21–23,25–28,34^ of the 24 meta-analyses; Fig. 5). We used three different methods that accounted for dependence between the effects coming from the same sample of participants: funnel asymmetry test (FAT)^46^ with the aggregates of the effect sizes of the same study, a three-parameter selection model (3PSM)^47^, and the combination of the proportion of statistical significance test (PSST) and the test of excess statistical significance (TESS^48^; for details, see Methods). We observed evidence of publication bias at least with one of the methods in 9^9,15,19,22,23,25–28^ of the 12 meta-analyses. That is, effect sizes of smaller primary studies tended to be larger (as suggested by FAT), positive and significant results were more likely to be published (i.e., 3PSM), or the proportion of positive results was higher than the expected proportion given the meta-analytic effect and variance (according to TESSPSST) in 9 of 12 meta-analyses. In general, the final effect was reduced after bias correction (Fig. 5), with the smallest values when the intercept of meta-regressive FAT models was used as a corrected estimation of the true effect (i.e., PET-PEESE method^49^): median of 0.01, 95% CI [−0.05, 0.05], that was non-significant in all the cases. However, 3PSM showed positive but reduced benefits in general (median of 0.16, 95% CI [0.12, 0.26]).

**Fig. 5.**
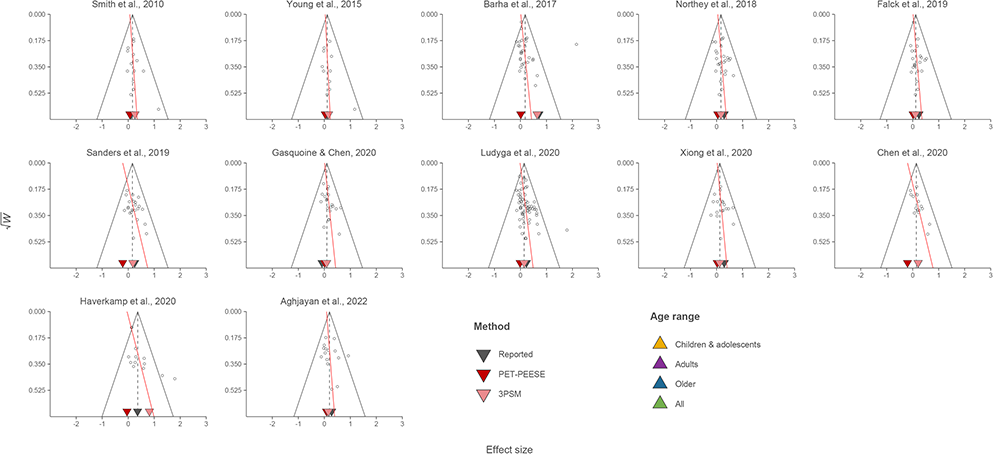
Assessment of publication bias across the meta-analyses included in the umbrella review. Funnel plot of 12 of the 24 meta-analyses. The aggregated outcomes of the primary studies are depicted with circles, whereas the red line represents the effect predicted by the PET-PEESE meta-regressive model, where a non-zero slope may indicate publication bias. The final effect reported in the original paper and the adjusted final effects are depicted with triangles at the bottom of each funnel plot. In general, there was evidence of publication bias suggesting that in most cases the benefits of physical activity on cognition were likely overestimated.

To further elucidate its impact on the field, we assessed publication bias in the entire set of primary studies. In addition to the three aforementioned classic methods to detect the presence of publication bias (i.e., FAT with aggregates, 3PSM, and TESSPSST), we also applied FAT using a multilevel model (multilevel FAT)^50^. All methods detected evidence of publication bias, suggesting the need of correcting the meta-analytic effect. The corrected effect was negligible with PET-PEESE procedure (multilevel PET-PEESE: *d* = −0.04, 95% CI [−0.16, 0.07], *p* = .420; PET-PEESE with aggregates: *d* = −0.01, 95% CI [−0.08, 0.06], *p* = .778). On the other hand, the adjusted effect increased with 3PSM, *d* = 0.31, 95% CI [0.15, 0.47], *p* < .001, although it also decreased with a larger set of cut points (one-tailed *p* values of .025, .05, .10, .20, .40, .50, and .70), *d* = −0.02, 95% CI [−0.34, 0.30], *p* = .898. Given the limitations of frequentist inferences about non-significant results, we applied robust Bayesian meta-analysis to obtain a model-averaged estimate of these two approaches of adjustment (i.e., selection models and PET-PEESE)^52^. Bayesian meta-analysis allows obtaining the best possible adjusted meta-analytic effect size among models with and without correction for publication bias. The model suggested strong evidence of publication bias, *BF*_pb_ = 648.91, although it led to inconclusive evidence of a positive effect, *BF*_10_ = 0.96, posterior mean estimate of *d* = 0.05, 95% CrI [−0.09, 0.14]. Therefore, besides the fast growth of this literature, the publication process seems to have favored the reporting of positive and significant results over null results, especially in small studies. These findings suggest that the true effect of exercise on cognition is probably smaller than originally reported in the meta-analyses and, despite the substantial number of individual studies, the available evidence is far from conclusive.

### Specification-curve analysis

As we described in the previous sections, the differences in the inclusion criteria of the reviewed meta-analyses may partly explain the discrepancies in their findings. However, even when including the same primary studies, their results might vary due to differences in the multiple preprocessing and analytic steps. The meta-analyses differed, for example, in the way they estimated the effect size and the variance, the use of influential moderators to adjust the outcome, the method for assessing publication bias, and the approach used to deal with the within-effects dependence. To examine the impact of all these decisions on the meta-analytic outcome, we conducted an exploratory (not pre-registered) specification-curve analysis^53^ with the entire set of primary studies we used in our full multilevel meta-analysis. The analysis showed that the final effect could vary greatly depending on preprocessing and analytic decisions (from *d* = ⎼0.05 to *d* = 0.47; Fig. 6). Some common specifications in the reviewed meta-analyses such as not dealing with within-study dependence using univariate models (9^7,8,14,19,27,30,31,33,34^ out of 24), not correcting for publication bias (14^7,8,14,15,18–20,23– 26,28,29,32^ out of 24) or doing so with a trim-and-fill method (2^9,34^ out of 24), which are not recommended in meta-analytic practice, led to higher effects and more likely to be significant. In contrast, some conservative decisions to increase the robustness of the outcomes (rarely adopted in the revised meta-analyses) reduced the final effect substantially: the use of multilevel models (4^17,21–23^ out of 24) and correcting for publication bias with PET-PEESE (2^9,22^ out of 24). Therefore, most of the meta-analyses opted for specifications that tend to find more positive and significant effects, whereas meta-analyses with more conservative decisions were underrepresented in the literature.

**Fig. 6.**
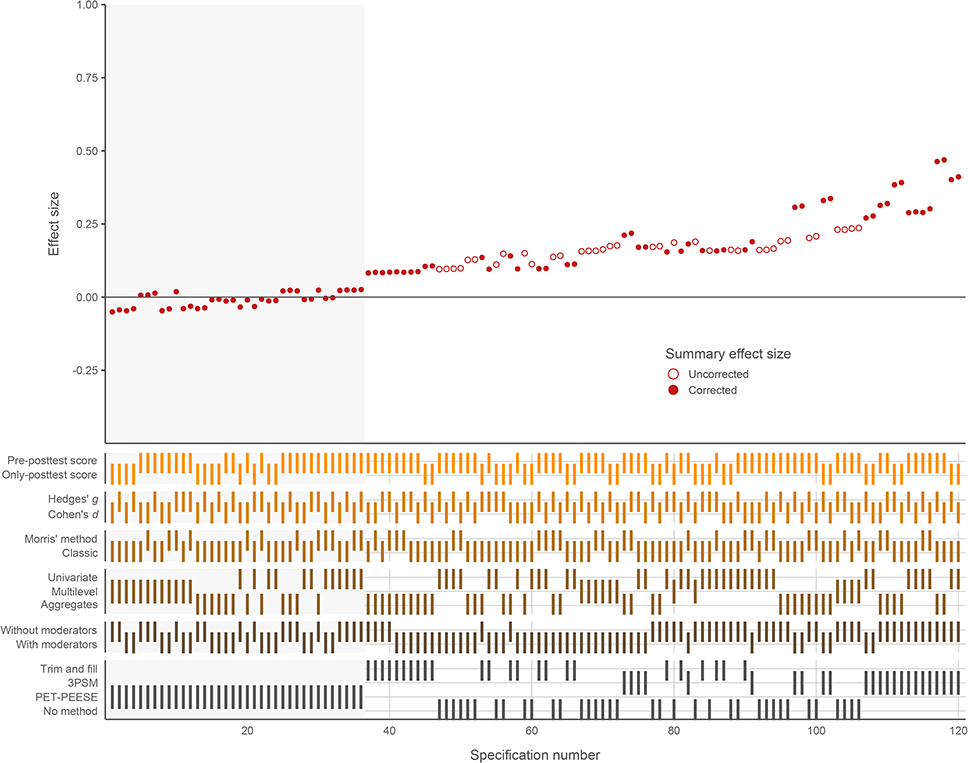
Specification curve of meta-analytic models. Summary effect size of the target studies (and its 95% confidence interval) varied across the multiple combinations of preprocessing and analytic decisions. Red empty effects represent the outcome of models without publication-bias adjustment, whereas red filled effects show the corrected summary effect. The light gray rectangular shade distinguishes non-significant results from significant ones. It is apparent the high variability of results and the disparity of conclusions that can be extracted from them. The present specification-curve analysis highlights the great impact of preprocessing and analytic decisions on the final outcome.

## Discussion

In this umbrella review, we examined the claim that regular physical exercise leads to cognitive gains across the lifespan. After reanalyzing 24 meta-analyses of RCTs, including a total of 109 primary studies and 11266 healthy participants, we found inconclusive evidence supporting the existence of a potential cognitive benefit derived from the regular practice of physical exercise in healthy populations. Our findings suggest that the effect of exercise on cognition reported in previous meta-analytic reviews has likely been overestimated and that, in general, the available causal evidence from RCTs on the exercise-cognition link is far from conclusive.

This review provides a fine-grained outline of the exponential growth of the exercise–cognition in healthy humans over the past fifty years. The rapid growth has provided insight into the potential benefits, risks, and pitfalls of implementing exercise-based interventions in the general population, resulting in a vast body of evidence. This evidence, often from underpowered RCTs and potentially biased meta-analyses, shows signs of unusual clustered-like growth (i.e., meta-analyses on the topic do not share primary sources of evidence), where the beneficial effect of physical exercise on cognition has been taken for granted despite the existence of several accounts that have shown mixed^12^ or contradictory findings^54^. In line with recent accounts^55^, we believe this exponential accumulation of low-quality evidence has led to stagnation rather than advance in the field hindering the discernment of the real existing effect.

Replication of published scientific findings is currently at the forefront of scientific debate. Since the publication of the seminal Reproducibility Project: Psychology^56^, there have been many initiatives in different research fields to test the extent to which the empirical evidence is solid and reliable^57–59^. The “*replication crisis*” has shaken the foundations of numerous fields and continuous to question many of the effects that have long been believed to be true. This umbrella review is part of the collective effort to promote transparency, openness, and reproducibility in science. Here, we delineate the structural weaknesses of the exercise–cognition meta-analytic literature, including the marked methodological, theoretical, and communicative issues. Below, we briefly develop a series of opportunities for improvement that may guide future studies in the field.

The exercise–cognition field has been flooded with individual experiments addressing this relationship on designs with low statistical power that yield estimates with low precision and stability. While conducting intervention studies with remarkably large sample sizes (see *Footnote 3*) is not within the reach of almost any individual laboratory (see Zotcheva et al.^60^ for a recent high-quality exception), simplification and standardization of experimental designs to increase statistical power and facilitate comparison of results and replication, proper active control groups, pre-registration, or multi-laboratory initiatives^61^ can definitely enhance the field.

Notably, epidemiological studies are an additional important source of evidence that suggests a potential role of physical exercise in cognitive improvement^62^. Nevertheless, such studies have also failed to show consistent evidence in support of the hypothesis that regular exercise boosts cognitive performance. While it is true that they have occasionally reported exercise-associated cognitive benefits, this has in some cases been explained by baseline differences^63^. In other cases, physical exercise was not associated with any cognitive advantage even after years of practice^43^. Future longitudinal studies could help to discern the existence of a positive impact of regular exercise on cognition, as well the specific role of different mediators and moderators.

In addition, it is important to cautiously reconsider the role of meta-analyses and the extent to which their results shed light on this particular topic^64,65^. Even though meta-analytic approaches have the potential to minimize some of the shortcomings of individual intervention studies^66^, their results largely depend on the quality of the included reports as well as on the methodological decisions followed to estimate a particular effect^67^. Thus, meta-analytic results do not necessarily represent the true effect of a particular phenomenon. As we show here, the particular conclusions from the different meta-analyses cannot be taken as the empirical evidence accumulated over years, but as selective slices of it. Moreover, subsampling all the available evidence (i.e., meta-analyzing a subset of primary studies) might lead to unreliable outcomes, likely departing from the true effect. It would also result in a reduced capacity to assess the impact of moderators and publication bias on the final result.

For conclusions of future meta-analyses to be translated into social recommendations and evidence-based policies, the reviews need to be as comprehensive as possible. This means that they should incorporate not only the knowledge from the primary literature, but also from previous reviews. Bearing in mind the marked publication bias in the exercise–cognition field, forthcoming meta-analyses should include gray literature contributing, predictably, with less optimistic results. Finally, at the preprocessing and analytic levels, there are many choices that are arguably preferred. For example, the use of an effect size estimate based on the pre-posttest changes to control for preexisting differences at baseline. We also encourage the use of multilevel models (e.g., RVE approach) accounting for the correlated structure of effects without averaging information and reducing heterogeneity by the identification of moderating variables in the model. Regarding the assessment of publication bias, factors such as the number of studies, heterogeneity, or the degree of publication bias present in the literature might affect the specific performance of the available methods^68^. A reasonable strategy is to identify the plausible conditions of the meta-analysis and interpret in consequence the findings. As an alternative, methods such as robust Bayesian meta-analysis allow weighting models regarding their fit to the evidence and obtaining one single meta-analytic effect size, without needing to choose among several outcomes. The exercise–cognition literature is currently big enough for the implementation of this type of analysis and to adjust in consequence the final effect.

The large number of published experiments and reviews on this topic sharply contrasts with the absence of a firm theoretical model of the mechanisms involved in exercise-induced cognitive improvements in humans is surprising. Research on animal models has certainly provided important insight into the possible neurobiological mechanisms. For example, it has been often reported in animals, and occasionally also in humans, that regular exercise 1) increases the release of key biochemical mediators of neuronal survival such as the brain-derived neurotrophic factor and other growth factors like the vascular endothelial growth factor or insulin-like growth factor type-1^69^; 2) counteracts grey and white matter atrophy as we age^70^; 3) increases vasculature, dendritic spine density and hippocampal complexity^71^; 4) enhances synaptic plasticity^72^; 5) increases resistance to brain insult^73^; 6) reduces central and peripheral inflammation^74^; and even 7) mobilizes gene expression profiles^75^. Despite the difficulty in extending these findings and the respective hypothetical mechanisms to humans (because of the inherent physiological and behavioral differences between humans and other species), certain concrete hypotheses have been proposed to explain the potential cognitive benefits of regular exercise in humans such as the cardiovascular hyphotesis^76^, the anti-inflammatory hyphotesis^74^ or the catecholamine hypothesis^77^. Although evidence-based, all these hypotheses adopt a reductionist approach by assuming that the mechanisms underlying exercise-related benefits stem merely from physiological processes at the molecular and cellular level, giving little relevance to the contextual complexity involved in the practice of physical exercise in humans. Indeed, physical exercise is much more than a way to trigger physiological changes. In fact, other theoretical models point to the cognitive and social enrichment accompanying physical exercise as the source of its putative cognitive benefits (i.e., the cognitive training hyphotesis^78^), or even the possibility that there are differences at the genetic level that might explain this association (i.e., the neuroselection hyphotesis^63^). Although there is data supporting each of these hypotheses, to date, none of them has been able to fully account for all the existing evidence which seems to point to a very complex relationship that is best described as a multifactorial phenomenon. This absence of consensus on a theoretical framework has been further accentuated by the myriad of experimental approaches that hamper the reconciliation of different findings^5^. Certainly, it is necessary to shift away from the metaphor of the brain as a muscle^79^ and further develop comprehensive theoretical models on the cognitive and neural mechanisms of these potential exercise-induced cognitive improvements.

The results reported here, together with the methodological and theoretical issues mentioned above, highlight the need to nuance claims regarding the potential cognitive benefits associated with the regular practice of physical exercise. However, the pressure for publishing is an endemic issue shared by researchers and mass media. In both cases, there is an urgency for publishing novel and eye-catching findings to attract public attention, which sometimes leads to oversimplification, misrepresentation, or overdramatization of scientific results without the nuances and limitations essential for proper interpretation. The exercise–cognition topic is not immune to this. Transparent practices throughout the research process (e.g., reporting bias-corrected effects) and accurate dissemination of scientific findings through the media would definitely improve the situation^80^. Nevertheless, this is a necessary but not sufficient step. Without the willingness of researchers to transparently report their results and databases, the collaborative efforts of editors to publish meaningful results (regardless of whether they are positive or not), and the commitment of the media to move away from hyped headlines and clickbait, it is a futile endeavor.

Although an umbrella review is considered to be the highest level of evidence in intervention research, it is important to acknowledge its limitations when interpreting the results. This review focuses exclusively on the impact of physical exercise, excluding exercise programs combined with any other cognitive intervention, on healthy populations. This leaves open the possibility that physical exercise may have a facilitating or protective effect on certain cognitive functions in individuals with certain diseases, such as Alzheimer’s disease, or that physical activities that include both physical and cognitive training, such as yoga, could enhance the benefits derived from physical exercise^81^. Additionally, this review only includes RCTs, but it is important to note that other sources of empirical evidence, such as observational or epidemiological studies, should also be considered. Lastly, the findings should be interpreted with caution as genetic and environmental factors may act as confounding variables.

In sum, our findings illustrate that current evidence from RCTs does not support a causal effect of regular physical exercise on cognitive enhancement, although it does not preclude it either. Consequently, and until stronger evidence accumulates, we urge caution in claims and recommendations linking exercise to cognitive benefits in the healthy population. Regarding current public health policies and guidelines for the promotion of physical exercise, we strongly believe there is no need to appeal to the alleged, as yet uncertain, cognitive benefits of physical exercise, especially when the current meta-analytic evidence from RCTs suggests that, even if the effect exists, it is notably small to assert its practical relevance (a Cohen’s *d* of 0.125, such as the adjusted effect found in the original meta-analysis by Ludyga et al.^9^, represents an explanation less than 1% of the variability between groups, or also corresponds with an increase of 2 IQ scores). The benefits of physical exercise on human well-being, especially with regard to physical health, are in themselves sufficient to justify evidence-based public health policies to promote its regular application in our daily lives^82^. Further, engaging in physical exercise brings not only physical but also social benefits, as we connect with others by forging social bonds and participating in collective activities that give us a sense of belonging. Further, engaging in physical exercise brings not only physical but also social benefits, as we connect with others by forging social bonds and participating in collective activities that give us a sense of belonging. Lastly, let us not forget the pleasure of doing something for its own sake. The value of exercising may lie simply in its enjoyable nature.

## Methods

### Literature search

We conducted a systematic literature search following the PRISMA guidelines^83^ (last search in July 2022) in Medline and Scopus using the following Boolean operators: *(“exercise” OR “physical activity” OR “physical exercise” OR “chronic exercise” OR “regular exercise”) AND (“cognition” OR “brain” OR “executive functions’’ OR “memory”) AND (metaanalysis OR meta-analysis)*. Additionally, we searched on Proquest and Google Scholar to identify unpublished meta-analyses meeting the inclusion criteria. Search was limited to papers published in English. The search was carried out independently by three authors (DH, DS, and LFC) who revised the titles and abstracts to identify possible additional publications. Subsequently, two authors (DH and LFC) revised the full text of these articles, and discrepancies between these authors were resolved by a third author (DS).

### Inclusion and exclusion criteria

We followed the Participant-Intervention-Comparison-Outcome (PICO) process to select the meta-analyses included in this umbrella review: (1) Participants: healthy participants of all ages and both sexes. Meta-analyses with clinical populations, including obesity or mild cognitive impairment, were excluded. However, if meta-analyses included healthy participants and this effect could be extracted separately, we considered only the effect of this population; (2) Intervention: RCTs studying the effects of a regular exercise program in any cognitive outcome with a minimum duration of two weeks and either involving aerobic exercise, resistance exercise, mixed exercises, or other physical activities (such as extracurricular physical activities); (3) Comparison: an active control group (in which participants completed a different exercise program or an alternative activity) or a passive control group (participants did not complete any exercise program); (4) Outcome: meta-analyses should report at least a measure of the global cognitive functioning or any specific cognitive domain (executive functions, attention, memory, etc.). To avoid confounding factors, we excluded meta-analyses or primary studies from reviewed meta-analyses involving mind-body, yoga, or exercise programs combined with any other intervention (e.g., cognitive training).

### Data extraction

The following data were extracted from the 274 primary studies included in the meta-analyses: (1) list of authors and year of publication; (2) list of authors and year of publication from each primary article included in the meta-analysis; (3) pooled number of participants for the experimental and control group; (4) sample age; (5) type of exercise intervention; (6) program duration; (7) cognitive outcome assessed; (8) type of control condition; (9) type and estimation method of effect size; (10) effect size, standard error, and confidence interval; (11) analysis of publication bias; (12) protocol registration; and (13) availability of data. Based on these data, we identified 109 primary studies that met the criteria of the present umbrella review (RCTs, healthy participants, physical exercise with no mind-body or cognitive training components, etc.). We then reviewed the specific inclusion and exclusion criteria of each meta-analysis and identified which of the primary studies meeting these criteria were available at the moment of their last reported search and should have therefore been included. When the meta-analyses did not report the date of the last search**^16,32^**or only reported the month**^14,15,19–21,25,27,29,33^**, we used the journal’s received date and the last day of the corresponding month as references, respectively. The final spreadsheet integrating all data is available at the following link: https://osf.io/e9zqf/.

Given the variety of experimental designs used in this literature, we decided to test the influence of the type of control (passive, control physical activity, or non-physical alternative activity), type of outcome (global cognition, executive functions, or other cognitive domains), and age range of the participants (children and adolescents, young adults, or older adults).

### Statistical analysis

To avoid the influence of using non-target exercise programs or samples with health conditions, we reanalyzed the effect sizes from the primary studies only with the target interventions and healthy populations. We used the standardized mean change between the pre and posttest scores with small-sample-bias correction as the estimator of the effect size in the main analyses. The pooled standard deviation was estimated as a combination of the pretest standard deviations of both groups^84^. For the main analyses, we estimated the variance using the formula

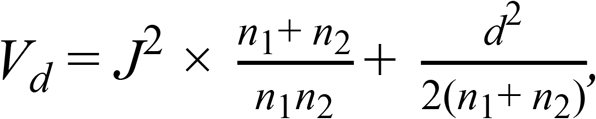

where *J* represents the correction factor of the small sample bias. Moreover, to prevent a disproportionate influence of those studies with an unbalanced number of participants in one group compared with the other (*n*_1_/*n*_2_ > 3/2 or < 2/3), the sampling variance was calculated by replacing the size of the greatest group with the size of the smallest one.

We performed the analysis for each meta-analysis using its set of primary studies and then implemented the model with the entire sample of studies. We used the robust variance estimation method^85^ (RVE) using the *robumeta*^86^ package for R statistical software to conduct multilevel models. This method allows dealing with a correlated structure of outcomes from the same primary study. To assess the influence of three moderating variables (type of control activity, type of cognitive outcome, and age range), we re-estimated the outcomes of the 24 meta-analyses for each level of the variables. In addition, we implemented separate RVE models for each moderator with the entire sample of studies. In the full model, we also tested the influence of pre-intervention performance as another factor that has explained part of the variability in the literature on cognitive training^80^. Even in RCTs, it is possible to observe baseline differences between small experimental and comparison groups due to sampling error. Because of a greater window of opportunity and/or regression toward the mean, the participants of the experimental group showed greater benefits when their performance at baseline was lower than the control group. We used the standardized mean difference of the pretest scores, *g*_pre_, as moderator.

We re-assessed publication bias in those reviewed meta-analyses with at least ten primary studies that met our inclusion criteria^8,9,14,15,19,21–23,25–28^. In our re-analysis, we accounted for dependence using aggregates of all the effect sizes from the same study included in each meta-analysis^87^ (i.e., averaging all effect sizes from the same sample of participants, thus reducing the number of outcomes per study to one). When the publication process favors the selective report of positive and significant results, underpowered studies (i.e., with smaller sample sizes and larger standard error) are more prone to show larger effects. Thus, this association between the size of the estimates and their precision can be assessed with a meta-regressive model that includes the standard error of the effect as a covariate^50^. A significant meta-regressive coefficient indicates a non-zero association (i.e., usually larger effects with smaller studies) and, thus, an asymmetrical distribution of effects, departing from a funnel shape. We tested for small-study bias (i.e., funnel asymmetry test or FAT) using a weighted-least squares meta-analysis with within-study aggregates. To prevent the artifactual dependence between the effect size and its precision estimate, we used an alternative formula to estimate the variance^88^: *W* = (*n*_1_ + *n*_2_) / (*n*_1 ×_ *n*_2_). This formula does not include the to-be-estimated effect size, as in the classic estimation of variance 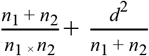, where the *d* represents the meta-analytic effect). In addition, we fitted a three-parameter selection model (3PSM)^47^ with aggregates. 3PSM is a version of the Vevea and Hedges’ selection model with only one cut point at the significance threshold (at *p*_one-tailed_ = .025) that estimates the probability of observing non-significant results over the significant ones (used as reference with a weight of 1). After estimating it, a likelihood-ratio test allows assessing if the adjusted model is a better fit. Assuming that positive and significant results would be more likely to be reported, 3PSM adjusts the final estimate and heterogeneity of a random-effects meta-analysis based on the *p* values of the individual effects. In addition, we used the proportion of statistical significance test (PSST) and the test of excess statistical significance (TESS) in combination^48^. PSST compares the observed proportion of significant studies with the theoretical proportion of studies that would report significant results given the conditions of the meta-analysis (i.e., fitted estimate, within-study variance, and heterogeneity). In TESS, the proportion of excess statistical significance is compared against the acceptable rate of false positives (i.e., 5% of type I errors). We implemented PET-PEESE^49^ and 3PSM^47^ with aggregates to adjust the final estimate for publication bias. The PET-PEESE procedure takes the intercept of the meta-regressive model as the best estimate of the underlying effect (i.e., the estimated effect when the sampling error is zero), whereas 3PSM corrects the effect size using the estimated probability that studies with certain *p* values were published. In some meta-analyses, 3PSM showed problems of convergence with a random-effects model but converged with a fixed-effects one^15,25,27^.

Subsequently, we examined publication bias with the entire sample of studies. To deal with correlated structure of outcomes (i.e., several effect sizes from the same sample of participants), we used an RVE multilevel model or aggregates from the same study^87^, regarding the method. FAT was implemented with aggregates (FAT with aggregates) and with an RVE multilevel model (FAT multilevel), as well as 3PSM, and the combination of PSST and TESS with study aggregates. To account for part of the heterogeneity, we included type of control and baseline performance as moderators in the model of the two FAT methods and 3PSM. In the case of PSST and TESS, we adjusted the observed effect sizes assuming active control and no baseline difference. Fitting a univariate meta-analysis with the aggregates, we used the coefficients of both moderators to adjust the observed effect (δ_obs_) assuming the use of active control group and no prior difference:

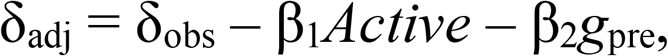

where β_1_ and β_2_ are the meta-regressive coefficients of the moderators, and *Active* is a dummy variable with the values 0 and 1 for active and passive control, respectively. We used PET-PEESE procedures and 3PSM to correct the final outcome for selective reporting and small-study bias. In addition, we conducted a robust Bayesian meta-analysis^52^ to reach a single model-average estimate across these two approaches of adjustment: selection models and PET-PEESE. For selection models, we used the six sets of p-value cut points defined in Bartoš et al.^52^: (1) two-sided selection, .05; (2) two-sided selection, .05 and .10; (3) one-sided, .05; (4) one-sided, .025 and .05; (5) one-sided, .05 and .50; (6) one-sided, .025, .05, and .50. Instead of selecting the result of one of the approaches, the Bayesian model weights the estimates of the different models (18 in total as we only conducted random-effects models) with the support they receive from the data. Two relevant advantages of this method are that robust Bayesian meta-analysis allows avoiding the choice between the outcome of one or other publication-bias approach, as well as it can distinguish between “absence of evidence” and “evidence of absence”. Based on previous meta-analyses and the literature that reflect a prior belief of a small effect of physical exercise, we selected a normal distribution centered at 0.35 and with one standard deviation as the prior for the effect in the alternative hypothesis. For the effect belonging to the null hypothesis, we assumed a normal distribution centered at 0 and equal standard deviation. We conducted the analysis only with random-effects models, that is assuming non-zero heterogeneity in all the cases, which is the most common scenario in psychological research.

Finally, we conducted an exploratory specification-curve analysis^53^ with all the primary studies. In the specification curve, we estimated the final effect size and its significance value for a total of 120 possible combinations of three preprocessing: (a) the standardized effect size was based on the pre-posttest change or only on the posttest difference between groups; (b) the use of Cohen’s *d* or Hedges’ *g*; and (c) the estimation of the variance of the effect size followed the classic formulation or the Morris’ proposal. For the specification curve, we selected these three preprocessing steps along with three additional analytic decision levels: (d) how to deal with within-study dependence (none strategy or assuming within-effects independence, an RVE multilevel model, and fitting a univariate model with aggregate effect sizes; (e) the inclusion or not of influential moderators to adjust the outcome (i.e., type of control group and baseline difference); and (f) the strategies to assess and correct the final outcome for publication bias (PET and PEESE methods, trim-and-fill approach, 3PSM, and no correction). The analysis led to 120 different combinations of specifications as the trim-and-fill method and 3PSM cannot be conducted with a multilevel model, and the Morris’ variance was only applied to pre-posttest estimates of the effect size (reducing the number of possible combinations from 192 to 120).

### Sample size estimation

We estimated that 652 participants are required to achieve a power of .80 for a small effect size such as *d* = 0.22 in a two-tailed two-sample *t*-test and an α of .05. This analysis corresponds with the classic between-group comparison in the pre-posttest change of the cognitive measure. However, another common analysis is to conduct a two-way mixed ANOVA with a between-group factor (i.e., treatment vs. control group) and moment as a within-participant factor (i.e., pretest vs. posttest). There, the key effect is the group-by-moment interaction, in which at least 654 participants are necessary to achieve an adequate power for a Cohen’s *f* = 0.11 (or, equivalently, an η^2^ = 0.012).

### Pre-registration

The methods and planned analyses of this umbrella review were pre-registered on June 9, 2020 at PROSPERO (CRD42020191357). The hypotheses and analysis plan were pre-registered in the OSF repository: https://osf.io/hfrpc. All deviations from the pre-registered procedures and analysis plans are transparently identified in the manuscript.

## Data availability

Data used to support the conclusions of this study are available at the OSF repository: https://osf.io/e9zqf/

## Code availability

Codes used for the analyses presented here are available at the OSF repository: https://osf.io/e9zqf/.

## Acknowledgements

This research was supported by a postdoctoral fellowship from the Junta de Andalucía to LFC (DOC00225), a predoctoral fellowship from the Spanish Ministry of Education, Culture and Sport (FPU17/02864) and a research grant from the Junta de Andalucía (PY20_00693) to RRC, a research grant from the Community of Madrid and the Rey Juan Carlos University (V-1159) to ALC, and two research grants from the Spanish Ministry of Economy and Competitiveness awarded to MAV (PID2020-118583GB-I00) and to DS (PID2019-105635GB-I00). The funders had no role in study design, data collection and analysis, decision to publish or preparation of the manuscript.

## Author Contributions

LFC, MAV, DH, ALC, PP and DS were involved in the original conceptualization. LFC, RRC, MAV, DH, ALC, PP and DS were responsible for developing the study methodology. LFC, DH and DS did the literature search. LFC, RRC, MAV, DH, ALC, PP and DS were responsible for data curation. RRC and MAV did the formal statistical analysis. LFC, RRC, DH and DS wrote the original draft. MAV, DH, ALC and PP edited and reviewed the manuscript.

## References

1. Sallis, J. F. et al. Physical activity in relation to urban environments in 14 cities worldwide: a cross-sectional study. The Lancet 387, 2207–2217 (2016).

2. Guiney, H. & Machado, L. Benefits of regular aerobic exercise for executive functioning in healthy populations. Psychonomic bulletin & review 20, 73–86 (2013).

3. Ding, D. et al. Physical activity guidelines 2020: comprehensive and inclusive recommendations to activate populations. The Lancet 396, 1780–1782 (2020).

4. Bull, F. C. et al. World Health Organization 2020 guidelines on physical activity and sedentary behaviour. British journal of sports medicine 54, 1451–1462 (2020).

5. Basso, J. C. & Suzuki, W. A. The Effects of Acute Exercise on Mood, Cognition, Neurophysiology, and Neurochemical Pathways: A Review. Brain Plasticity 2, 127–152 (2017).

6. Tomporowski, P. D. & Pesce, C. Exercise, sports, and performance arts benefit cognition via a common process. Psychological bulletin 145, 929 (2019).

7. Angevaren, M., Aufdemkampe, G., Verhaar, H., Aleman, A. & Vanhees, L. Physical activity and enhanced fitness to improve cognitive function in older people without known cognitive impairment. in Cochrane Database of Systematic Reviews (ed. The Cochrane Collaboration) CD005381.pub3 (John Wiley & Sons, Ltd, 2008). doi:10.1002/14651858.CD005381.pub3.

8. Young, J., Angevaren, M., Rusted, J. & Tabet, N. Aerobic exercise to improve cognitive function in older people without known cognitive impairment. Cochrane Database of Systematic Reviews (2015).

9. Ludyga, S., Gerber, M., Pühse, U., Looser, V. N. & Kamijo, K. Systematic review and meta-analysis investigating moderators of long-term effects of exercise on cognition in healthy individuals. Nature human behaviour 4, 603–612 (2020).

10. Verburgh, L., Konigs, M., Scherder, E. J. A. & Oosterlaan, J. Physical exercise and executive functions in preadolescent children, adolescents and young adults: a meta-analysis. British Journal of Sports Medicine 48, 973–979 (2014).

11. Singh, A. S. et al. Effects of physical activity interventions on cognitive and academic performance in children and adolescents: a novel combination of a systematic review and recommendations from an expert panel. British journal of sports medicine 53, 640–647 (2019).

12. Diamond, A. & Ling, D. S. Aerobic-Exercise and resistance-training interventions have been among the least effective ways to improve executive functions of any method tried thus far. (2019).

13. Erickson, K. I. et al. Physical Activity, Cognition, and Brain Outcomes: A Review of the 2018 Physical Activity Guidelines. Med Sci Sports Exerc 51, 1242–1251 (2019).

14. Colcombe, S. & Kramer, A. F. Fitness effects on the cognitive function of older adults a meta-analytic study. Psychological science 14, 125–130 (2003).

15. Smith, P. J. et al. Aerobic Exercise and Neurocognitive Performance: A Meta-Analytic Review of Randomized Controlled Trials: Psychosomatic Medicine 72, 239–252 (2010).

16. Scherder, E. et al. Executive functions of sedentary elderly may benefit from walking: a systematic review and meta-analysis. The American Journal of Geriatric Psychiatry 22, 782–791 (2014).

17. Lindheimer, J. B., O’Connor, P. J. & Dishman, R. K. Quantifying the placebo effect in psychological outcomes of exercise training: a meta-analysis of randomized trials. Sports Medicine 45, 693–711 (2015).

18. Jackson, W. M., Davis, N., Sands, S. A., Whittington, R. A. & Sun, L. S. Physical activity and cognitive development: a meta-analysis. Journal of neurosurgical anesthesiology 28, 373–380 (2016).

19. Barha, C. K., Davis, J. C., Falck, R. S., Nagamatsu, L. S. & Liu-Ambrose, T. Sex differences in exercise efficacy to improve cognition: A systematic review and meta-analysis of randomized controlled trials in older humans. Frontiers in Neuroendocrinology 46, 71–85 (2017).

20. Rathore, A. & Lom, B. The effects of chronic and acute physical activity on working memory performance in healthy participants: a systematic review with meta-analysis of randomized controlled trials. Systematic reviews 6, 1–16 (2017).

21. Northey, J. M., Cherbuin, N., Pumpa, K. L., Smee, D. J. & Rattray, B. Exercise interventions for cognitive function in adults older than 50: a systematic review with meta-analysis. Br J Sports Med 52, 154–160 (2018).

22. Falck, R. S., Davis, J. C., Best, J. R., Crockett, R. A. & Liu-Ambrose, T. Impact of exercise training on physical and cognitive function among older adults: a systematic review and meta-analysis. Neurobiology of aging 79, 119–130 (2019).

23. Sanders, L. M., Hortobagyi, T., la Bastide-van Gemert, S., van der Zee, E. A. & van Heuvelen, M. J. Dose-response relationship between exercise and cognitive function in older adults with and without cognitive impairment: a systematic review and meta-analysis. PloS one 14, e0210036 (2019).

24. Xue, Y., Yang, Y. & Huang, T. Effects of chronic exercise interventions on executive function among children and adolescents: a systematic review with meta-analysis. Br J Sports Med 53, 1397–1404 (2019).

25. Gasquoine, P. G. & Chen, P.-Y. Effect of physical exercise on popular measures of executive function in older, nonclinical, participants of randomized controlled trials: A meta-analytic review. Applied Neuropsychology: Adult 1–9 (2020).

26. Xiong, J., Ye, M., Wang, L. & Zheng, G. Effects of physical exercise on executive function in cognitively healthy older adults: A systematic review and meta-analysis of randomized controlled trials. International Journal of Nursing Studies 114, 103810 (2021).

27. Chen, F.-T. et al. Effects of Exercise Training Interventions on Executive Function in Older Adults: A Systematic Review and Meta-Analysis. Sports Med 50, 1451–1467 (2020).

28. Haverkamp, B. F. et al. Effects of physical activity interventions on cognitive outcomes and academic performance in adolescents and young adults: A meta-analysis. Journal of Sports Sciences 38, 2637–2660 (2020).

29. Hoffmann, C. M., Petrov, M. E. & Lee, R. E. Aerobic physical activity to improve memory and executive function in sedentary adults without cognitive impairment: A systematic review and meta-analysis. Preventive Medicine Reports 23, 101496 (2021).

30. Zhidong, C., Wang, X., Yin, J., Song, D. & Chen, Z. Effects of physical exercise on working memory in older adults: a systematic and meta-analytic review. European Review of Aging and Physical Activity 18, 1–15 (2021).

31. Amatriain-Fernández, S., Ezquerro García-Noblejas, M. & Budde, H. Effects of chronic exercise on the inhibitory control of children and adolescents: A systematic review and meta-analysis. Scandinavian Journal of Medicine & Science in Sports 31, 1196–1208 (2021).

32. Meli, A. M., Ali, A., Mhd Jalil, A. M., Mohd Yusof, H. & Tan, M. M. Effects of Physical Activity and Micronutrients on Cognitive Performance in Children Aged 6 to 11 Years: A Systematic Review and Meta-Analysis of Randomized Controlled Trials. Medicina 58, 57 (2021).

33. Zhao, Y. et al. Physical Activity and Cognition in Sedentary Older Adults: A Systematic Review and Meta-Analysis. Journal of Alzheimer’s Disease 1–12 (2022).

34. Aghjayan, S. L. et al. Aerobic exercise improves episodic memory in late adulthood: a systematic review and meta-analysis. Communications medicine 2, 1–11 (2022).

35. Colcombe, S. J. et al. Cardiovascular fitness, cortical plasticity, and aging. Proceedings of the National Academy of Sciences of the United States of America 101, 3316–3321 (2004).

36. Lakes, K. D. & Hoyt, W. T. Promoting self-regulation through school-based martial arts training. Journal of Applied Developmental Psychology 25, 283–302 (2004).

37. Muscari, A. et al. Chronic endurance exercise training prevents aging-related cognitive decline in healthy older adults: a randomized controlled trial. International journal of geriatric psychiatry 25, 1055–1064 (2010).

38. Voss, M. W. et al. Plasticity of brain networks in a randomized intervention trial of exercise training in older adults. Frontiers in aging neuroscience 2, 32 (2010).

39. Voss, M. W. et al. The influence of aerobic fitness on cerebral white matter integrity and cognitive function in older adults: Results of a one-year exercise intervention. Human brain mapping 34, 2972–2985 (2013).

40. Vidoni, E. D. et al. Effect of aerobic exercise on amyloid accumulation in preclinical Alzheimer’s: a 1-year randomized controlled trial. PloS one 16, e0244893 (2021).

41. León, J., Ureña, A., Bolaños, M. J., Bilbao, A. & Oña, A. A combination of physical and cognitive exercise improves reaction time in persons 61-84 years old. Journal of aging and physical activity 23, 72–77 (2015).

42. Bouaziz, W. et al. Effects of a short-term Interval Aerobic Training Programme with active Recovery bouts (IATP-R) on cognitive and mental health, functional performance and quality of life: A randomised controlled trial in sedentary seniors. International journal of clinical practice 73, e13219 (2019).

43. Sink, K. M. et al. Effect of a 24-month physical activity intervention vs health education on cognitive outcomes in sedentary older adults: the LIFE randomized trial. Jama 314, 781–790 (2015).

44. Lind, R. R. et al. Improved cognitive performance in preadolescent Danish children after the school-based physical activity programme “FIFA 11 for Health” for Europe–A cluster-randomised controlled trial. European Journal of Sport Science 18, 130–139 (2018).

45. Ansai, J. H. & Rebelatto, J. R. Effect of two physical exercise protocols on cognition and depressive symptoms in oldest-old people: A randomized controlled trial. Geriatrics & gerontology international 15, 1127–1134 (2015).

46. Sterne, J. A. & Egger, M. Funnel plots for detecting bias in meta-analysis: guidelines on choice of axis. Journal of clinical epidemiology 54, 1046–1055 (2001).

47. Vevea, J. L. & Hedges, L. V. A general linear model for estimating effect size in the presence of publication bias. Psychometrika 60, 419–435 (1995).

48. Stanley, T. D., Doucouliagos, H., Ioannidis, J. P. & Carter, E. C. Detecting publication selection bias through excess statistical significance. Research Synthesis Methods 12, 776–795 (2021).

49. Stanley, T. D. & Doucouliagos, H. Meta-regression approximations to reduce publication selection bias. Research Synthesis Methods 5, 60–78 (2014).

50. Egger, M., Smith, G. D., Schneider, M. & Minder, C. Bias in meta-analysis detected by a simple, graphical test. Bmj 315, 629–634 (1997).

51. Rodgers, M. A. & Pustejovsky, J. E. Evaluating meta-analytic methods to detect selective reporting in the presence of dependent effect sizes. Psychological methods 26, 141 (2021).

52. Bartoš, F., Maier, M., Wagenmakers, E.-J., Doucouliagos, H. & Stanley, T. D. Robust Bayesian Meta-Analysis: Model-Averaging Across Complementary Publication Bias Adjustment Methods. Research Synthesis Methods (2021).

53. Simonsohn, U., Simmons, J. P. & Nelson, L. D. Specification curve analysis. Nature Human Behaviour 4, 1208–1214 (2020).

54. Coen, R. F., Lawlor, B. A. & Kenny, R. Failure to demonstrate that memory improvement is due either to aerobic exercise or increased hippocampal volume. Proceedings of the National Academy of Sciences 108, E89–E89 (2011).

55. Chu, J. S. & Evans, J. A. Slowed canonical progress in large fields of science. Proceedings of the National Academy of Sciences 118, (2021).

56. Collaboration, O. S. Estimating the reproducibility of psychological science. Science 349, aac4716 (2015).

57. Errington, T. M. et al. Investigating the replicability of preclinical cancer biology. Elife 10, e71601 (2021).

58. Camerer, C. F. et al. Evaluating replicability of laboratory experiments in economics. Science 351, 1433–1436 (2016).

59. Camerer, C. F. et al. Evaluating the replicability of social science experiments in Nature and Science between 2010 and 2015. Nature Human Behaviour 2, 637–644 (2018).

60. Zotcheva, E. et al. Effects of 5 Years Aerobic Exercise on Cognition in Older Adults: The Generation 100 Study: A Randomized Controlled Trial. Sports Medicine 52, 1689–1699 (2022).

61. Kvarven, A., Strømland, E. & Johannesson, M. Comparing meta-analyses and preregistered multiple-laboratory replication projects. Nature Human Behaviour 4, 423–434 (2020).

62. Rockwood, K. & Middleton, L. Physical activity and the maintenance of cognitive function. Alzheimer’s & dementia 3, S38–S44 (2007).

63. Belsky, D. W. et al. Cardiorespiratory fitness and cognitive function in midlife: Neuroprotection or neuroselection?: Fitness and Cognitive Function. Annals of Neurology 77, 607–617 (2015).

64. Lau, J., Ioannidis, J. P. & Schmid, C. H. Summing up evidence: one answer is not always enough. The lancet 351, 123–127 (1998).

65. Pogue, J. & Yusuf, S. Overcoming the limitations of current meta-analysis of randomised controlled trials. The Lancet 351, 47–52 (1998).

66. Hunter, J. E. & Schmidt, F. L. Methods of meta-analysis: Correcting error and bias in research findings. (Sage, 2004).

67. Simonsohn, U., Simmons, J. & Nelson, L. D. Above averaging in literature reviews. Nature Reviews Psychology 1–2 (2022).

68. Carter, E. C., Schönbrodt, F. D., Gervais, W. M. & Hilgard, J. Correcting for bias in psychology: A comparison of meta-analytic methods. Advances in Methods and Practices in Psychological Science 2, 115–144 (2019).

69. Maass, A. et al. Relationships of peripheral IGF-1, VEGF and BDNF levels to exercise-related changes in memory, hippocampal perfusion and volumes in older adults. Neuroimage 131, 142–154 (2016).

70. Gow, A. J. et al. Neuroprotective lifestyles and the aging brain: activity, atrophy, and white matter integrity. Neurology 79, 1802–1808 (2012).

71. Stranahan, A. M. et al. Voluntary exercise and caloric restriction enhance hippocampal dendritic spine density and BDNF levels in diabetic mice. Hippocampus 19, 951–961 (2009).

72. Christie, B. R. et al. Exercising our brains: how physical activity impacts synaptic plasticity in the dentate gyrus. Neuromolecular medicine 10, 47–58 (2008).

73. Stummer, W., Weber, K., Tranmer, B., Baethmann, A. & Kempski, O. Reduced mortality and brain damage after locomotor activity in gerbil forebrain ischemia. Stroke 25, 1862–1869 (1994).

74. Cotman, C. W., Berchtold, N. C. & Christie, L.-A. Exercise builds brain health: key roles of growth factor cascades and inflammation. Trends in Neurosciences 30, 464–472 (2007).

75. Farmer, J. et al. Effects of voluntary exercise on synaptic plasticity and gene expression in the dentate gyrus of adult male Sprague–Dawley rats in vivo. Neuroscience 124, 71–79 (2004).

76. Tarumi, T. & Zhang, R. The role of exercise-induced cardiovascular adaptation in brain health. Exercise and sport sciences reviews 43, 181–189 (2015).

77. Exercise-Cognition Interaction. (Academic Press, 2016).

78. Mann, D. T., Williams, A. M., Ward, P. & Janelle, C. M. Perceptual-cognitive expertise in sport: A meta-analysis. Journal of Sport and Exercise Psychology 29, 457–478 (2007).

79. Simons, D. J. et al. Do “brain-training” programs work? Psychological Science in the Public Interest 17, 103–186 (2016).

80. Román-Caballero, R., Sanabria, D. & Ciria, L. F. Let’s go beyond “the effect of” physical exercise and musical training on cognition: much more than a significant effect. PsyArXiv (2021).

81. Gavelin, H. M. et al. Combined physical and cognitive training for older adults with and without cognitive impairment: A systematic review and network meta-analysis of randomized controlled trials. Ageing research reviews 66, 101232 (2021).

82. Ekkekakis, P. Why Is Exercise Underutilized in Clinical Practice Despite Evidence It Is Effective? Lessons in Pragmatism From the Inclusion of Exercise in Guidelines for the Treatment of Depression in the British National Health Service. Kinesiology Review 10, 29–50 (2020).

83. Moher, D., Liberati, A., Tetzlaff, J., Altman, D. G. & Group, T. P. Preferred Reporting Items for Systematic Reviews and Meta-Analyses: The PRISMA Statement. PLOS Medicine 6, e1000097 (2009).

84. Morris, S. B. Estimating effect sizes from pretest-posttest-control group designs. Organizational research methods 11, 364–386 (2008).

85. Hedges, L. V., Tipton, E. & Johnson, M. C. Robust variance estimation in meta-regression with dependent effect size estimates. Research synthesis methods 1, 39–65 (2010).

86. Fisher, Z., Tipton, E., Zhipeng, H. & Fisher, M. Z. Package ‘robumeta’. (2017).

87. Borenstein, M., Hedges, L. V., Higgins, J. P. & Rothstein, H. R. Introduction to meta-analysis. (John Wiley & Sons, 2021).

88. Pustejovsky, J. E. & Rodgers, M. A. Testing for funnel plot asymmetry of standardized mean differences. Research Synthesis Methods 10, 57–71 (2019).

