## Supplementary material for "An umbrella review of randomized control trials on the effects of physical exercise on cognition"

### Methodological quality evaluation

The quality of the meta-analyses included in this review was assessed using the validated scale Assessing the Methodology Quality of Systematic Review 2 (AMSTAR 2) checklist. The AMSTAR 2 scale is developed to assess the quality of systematic reviews and meta-analyses that includes both randomized and non-randomized controlled trials. The scale is composed of 16 items, assessing questions regarding the use of the PICO criteria, registration of the review protocol, assessment of the risk of bias in the literature, adequate analysis of the data, analysis publication bias, etc. Each item can be scored as “yes” and “no”, if they fulfill several criteria. Although a final score might be calculated, we followed the authors' recommendation of assessing the quality of the reviews based on critical domains rather than by a final score (Shea et al., 2017). Nonetheless, we provide the percentage of articles that fulfill each criterion. Additionally, in our pre-registration form, we stated that we would also assess the quality of the included articles with another scale (Grading of Recommendations Assessment, Development and Evaluation (Guyatt et al., 2008), but given that many of the items overlap with the AMSTAR 2 scale, we decided to omit it. The results of the critical domains assessed in the AMSTAR 2 scale are available here: *link temporarily removed as part of Double Blind Peer Review process*.

### Reasons of Primary Study Exclusion

#### A. With Some Health Condition or Cognitive Impairment

Babaei, P., Azali Alamdari, K., Soltani Tehrani, B., & Damirchi, A. (2013). Effect of six weeks of endurance exercise and following detraining on serum brain derived neurotrophic factor and memory performance in middle aged males with metabolic syndrome. *Journal of Sports Medicine and Physical Fitness, 53*(4), 437–443.

Bae, S., Lee, S., Lee, S., Jung, S., Makino, K., Harada, K., ... & Shimada, H. (2019). The effect of a multicomponent intervention to promote community activity on cognitive function in older adults with mild cognitive impairment: a randomized controlled trial. *Complementary Therapies in Medicine, 42*, 164–169. https://doi.org/10.1016/j.ctim.2018.11.011

Baker, L. D., Frank, L. L., Foster-Schubert, K., Green, P. S., Wilkinson, C. W., McTiernan, A., ... & Craft, S. (2010). Effects of aerobic exercise on mild cognitive impairment: a controlled trial. *Archives of Neurology, 67*(1), 71–79. https://doi.org/10.1001/archneurol.2009.307

Barnes, D. E., Santos-Modesitt, W., Poelke, G., Kramer, A. F., Castro, C., Middleton, L. E., & Yaffe, K. (2013). The Mental Activity and eXercise (MAX) trial: a randomized controlled trial to enhance cognitive function in older adults. *JAMA Internal Medicine, 173*(9), 797–804. https://doi.org/10.1001/jamainternmed.2013.189

Blumenthal, J. A., Smith, P. J., Mabe, S., Hinderliter, A., Welsh-Bohmer, K., Browndyke, J. N., ... & Sherwood, A. (2020). Longer Term Effects of Diet and Exercise on Neurocognition: 1-Year Follow-up of the ENLIGHTEN Trial. *Journal of the American Geriatrics Society, 68*(3), 559–568. https://doi.org/10.1111/jgs.16252

Bolandzadeh, N., Tam, R., Handy, T. C., Nagamatsu, L. S., Hsu, C. L., Davis, J. C., ... & Liu-Ambrose, T. (2015). Resistance training and white matter lesion progression in older women: exploratory analysis of a 12-month randomized controlled trial. *Journal of the American Geriatrics Society, 63*(10), 2052–2060. https://doi.org/10.1111/jgs.13644

Bossers, W. J., van der Woude, L. H., Boersma, F., Hortobágyi, T., Scherder, E. J., & van Heuvelen, M. J. (2015). A 9-week aerobic and strength training program improves cognitive and motor function in patients with dementia: a randomized, controlled trial. *The American Journal of Geriatric Psychiatry, 23*(11), 1106–1116. https://doi.org/10.1016/j.jagp.2014.12.191

Busse, A. L., Jacob Filho, W., Magaldi, R. M., Coelho, V. A., Melo, A. C., & Betoni, R. A. (2008). Effects of resistance training exercise on cognitive performance in elderly individuals with memory impairment: results of a controlled trial. *Einstein, 6*(4), 402–407.

Chen, S.-R., Tseng, C.-L., Kuo, S.-Y., & Chang, Y.-K. (2016). Effects of a physical activity intervention on autonomic and executive functions in obese young adolescents: A randomized controlled trial. *Health Psychology, 35*(10), 1120–1125. https://doi.org/10.1037/hea0000390

Crova, C., Struzzolino, I., Marchetti, R., Masci, I., Vannozzi, G., Forte, R., & Pesce, C. (2014). Cognitively challenging physical activity benefits executive function in overweight children. *Journal of Sports Sciences, 32*(3), 201–211. https://doi.org/10.1080/02640414.2013.828849

Daley, A. J., Crank, H., Saxton, J. M., Mutrie, N., Coleman, R., & Roalfe, A. (2007). Randomized trial of exercise therapy in women treated for breast cancer. *Journal of Clinical Oncology, 25*(13), 1713–1721. doi: 10.1200/JCO.2006.09.5083

Damirchi, A., Hosseini, F., & Babaei, P. (2018). Mental training enhances cognitive function and BDNF more than either physical or combined training in elderly women with MCI: a small-scale study. *American Journal of Alzheimer's Disease & Other Dementias, 33*(1), 20–29. https://doi.org/10.1177/1533317517727068

Davis, J. C., Bryan, S., Marra, C. A., Sharma, D., Chan, A., Beattie, B. L., ... & Liu-Ambrose, T. (2013). An economic evaluation of resistance training and aerobic training versus balance and toning exercises in older adults with mild cognitive impairment. *PloS ONE, 8*(5), e63031. https://doi.org/10.1371/journal.pone.0063031

Davis, C. L., Tomporowski, P. D., Boyle, C. A., Waller, J. L., Miller, P. H., Naglieri, J. A., & Gregoski, M. (2007). Effects of aerobic exercise on overweight children's cognitive functioning: a randomized controlled trial. Research Quarterly for Exercise and Sport, 78(5), 510–519.

Davis, C. L., Tomporowski, P. D., McDowell, J. E., Austin, B. P., Miller, P. H., Yanasak, N. E., ... & Naglieri, J. A. (2011). Exercise improves executive function and achievement and alters brain activation in overweight children: a randomized, controlled trial. *Health Psychology, 30*(1), 91–98. https://doi.org/10.1037/a0021766

de Souto Barreto, P., Cesari, M., Denormandie, P., Armaingaud, D., Vellas, B., & Rolland, Y. (2017). Exercise or social intervention for nursing home residents with dementia: a pilot randomized, controlled trial. *Journal of the American Geriatrics Society, 65*(9), E123-E129. https://doi.org/10.1111/jgs.14947

Dillon, K., & Prapavessis, H. (2020). REducing SEDENTary behavior among mild to moderate cognitively impaired assisted living residents: a pilot randomized controlled trial (RESEDENT study). *Journal of Aging and Physical Activity, 29*(1), 27–35. https://doi.org/10.1123/japa.2019-0440

Doi, T., Verghese, J., Makizako, H., Tsutsumimoto, K., Hotta, R., Nakakubo, S., ... & Shimada, H. (2017). Effects of cognitive leisure activity on cognition in mild cognitive impairment: results of a randomized controlled trial. *Journal of the American Medical Directors Association, 18*(8), 686–691. https://doi.org/10.1016/j.jamda.2017.02.013

Donnezan, L. C., Perrot, A., Belleville, S., Bloch, F., & Kemoun, G. (2018). Effects of simultaneous aerobic and cognitive training on executive functions, cardiovascular fitness and functional abilities in older adults with mild cognitive impairment. *Mental Health and Physical Activity, 15*, 78–87. https://doi.org/10.1016/j.mhpa.2018.06.001

Eggermont, L. H. P., Swaab, D. F., Hol, E. M., & Scherder, E. J. A. (2009). Walking the line: a randomised trial on the effects of a short term walking programme on cognition in dementia. *Journal of Neurology, Neurosurgery & Psychiatry, 80*(7), 802–804. http://dx.doi.org/10.1136/jnnp.2008.158444

Emery, C. F., Schein, R. L., Hauck, E. R., & MacIntyre, N. R. (1998). Psychological and cognitive outcomes of a randomized trial of exercise among patients with chronic obstructive pulmonary disease. *Health Psychology, 17*(3), 232–240. https://doi.org/10.1037/0278-6133.17.3.232

Fiatarone Singh, M. A., Gates, N., Saigal, N., Wilson, G. C., Meiklejohn, J., Brodaty, H., ... & Valenzuela, M. (2014). The Study of Mental and Resistance Training

(SMART) study—resistance training and/or cognitive training in mild cognitive impairment: a randomized, double-blind, double-sham controlled trial. *Journal of the American Medical Directors Association, 15*(12), 873–880.

Gaitán, J. M., Boots, E. A., Dougherty, R. J., Oh, J. M., Ma, Y., Edwards, D. F., ... & Okonkwo, O. C. (2019). Brain glucose metabolism, cognition, and cardiorespiratory fitness following exercise training in adults at risk for Alzheimer's disease. *Brain Plasticity, 5*(1), 83–95. doi: 10.3233/BPL-190093

Gallotta, M. C., Emerenziani, G. P., Iazzoni, S., Meucci, M., Baldari, C., & Guidetti, L. (2015). Impacts of coordinative training on normal weight and overweight/obese children's attentional performance. *Frontiers in Human Neuroscience, 9*, 577. https://doi.org/10.3389/fnhum.2015.00577

Ghahramani, M. H., Sohrabi, M., & Besharat, M. A. (2016). The effects of physical activity on impulse control, attention, decision-making and motor functions in students with high and low impulsivity. *Biosciences Biotechnology Research Asia, 13*(3), 1689–1696. http://dx.doi.org/10.13005/bbra/2318

Gothe, N. P., Kramer, A. F., & McAuley, E. (2014). The effects of an 8-week Hatha yoga intervention on executive function in older adults. *Journals of Gerontology Series A: Biomedical Sciences and Medical Sciences, 69*(9), 1109–1116. https://doi.org/10.1093/gerona/glu095

Gschwind, Y. J., Eichberg, S., Ejupi, A., de Rosario, H., Kroll, M., Marston, H. R., ... & Delbaere, K. (2015). ICT-based system to predict and prevent falls (iStoppFalls): results from an international multicenter randomized controlled trial. *European Review of Aging and Physical Activity, 12*(1), 1–11. https://doi.org/10.1186/s11556-015-0155-6

Hariprasad, V. R., Koparde, V., Sivakumar, P. T., Varambally, S., Thirthalli, J., Varghese, M., ... & Gangadhar, B. N. (2013). Randomized clinical trial of yoga-based intervention in residents from elderly homes: Effects on cognitive function. Indian Journal of Psychiatry, 55(Suppl 3), S357–S363. https://doi.org/10.4103/0019-5545.116308.

Hars, M., Herrmann, F. R., Gold, G., Rizzoli, R., & Trombetti, A. (2014). Effect of music-based multitask training on cognition and mood in older adults. *Age and Ageing, 43*(2), 196–200. https://doi.org/10.1093/ageing/aft163

Hiyama, Y., Yamada, M., Kitagawa, A., Tei, N., & Okada, S. (2012). A four-week walking exercise programme in patients with knee osteoarthritis improves the ability of dual-task performance: a randomized controlled trial. *Clinical Rehabilitation, 26*(5), 403–412. https://doi.org/10.1177/0269215511421028

Hoffman, B. M., Blumenthal, J. A., Babyak, M. A., Smith, P. J., Rogers, S. D., Doraiswamy, P. M., & Sherwood, A. (2008). Exercise fails to improve neurocognition in depressed middle-aged and older adults. *Medicine and Science in Sports and Exercise, 40*(7), 1344–1352. https://doi.org/10.1249/mss.0b013e31816b877c

Hsu, C.L., Best, J.R., Wang, S., Voss, M.W., Hsiung, R.G., Munkacsy, M., Cheung, W., Handy, T.C., Liu-Ambrose, T., 2017. The impact of aerobic exercise on fronto-parietal network connectivity and its relation to mobility: an exploratory analysis of a 6-month randomized controlled trial. *Frontiers in Human Neuroscience, 11*, 344. https://doi.org/10.3389/fnhum.2017.00344

Hu, J. P., Guo, Y. H., Wang, F., Zhao, X. P., Zhang, Q. H., & Song, Q. H. (2014). Exercise improves cognitive function in aging patients. International Journal of Clinical and Experimental Medicine, 7(10), 3144–3149.

Huang, T., Larsen, K. T., Jepsen, J. R. M., Møller, N. C., Thorsen, A. K., Mortensen, E. L., & Andersen, L. B. (2015). Effects of an obesity intervention program on cognitive function in children: A randomized controlled trial. *Obesity, 23*(10), 2101–2108. https://doi.org/10.1002/oby.21209

Kamegaya, T., Araki, Y., Kigure, H., Long-Term-Care Prevention Team of Maebashi City, & Yamaguchi, H. (2014). Twelve-week physical and leisure activity programme improved cognitive function in community-dwelling elderly subjects: a randomized controlled trial. *Psychogeriatrics, 14*(1), 47–54. https://doi.org/10.1111/psyg.12038

Kemoun, G., Thibaud, M., Roumagne, N., Carette, P., Albinet, C., Toussaint, L., ... & Dugué, B. (2010). Effects of a physical training programme on cognitive function and walking efficiency in elderly persons with dementia. *Dementia and Geriatric Cognitive Disorders, 29*(2), 109–114. https://doi.org/10.1159/000272435

Khatri, P., Blumenthal, J. A., Babyak, M. A., Craighead, W. E., Herman, S., Baldewicz, T., ... & Krishnan, K. R. (2001). Effects of exercise training on cognitive functioning among depressed older men and women. Journal of Aging and Physical Activity, 9(1), 43–57. https://doi.org/10.1123/japa.9.1.43

Krafft, C. E., Schwarz, N. F., Chi, L., Weinberger, A. L., Schaeffer, D. J., Pierce, J. E., ... & McDowell, J. E. (2014). An 8-month randomized controlled exercise trial alters brain activation during cognitive tasks in overweight children. *Obesity, 22*(1), 232–242. https://doi.org/10.1002/oby.20518

Kwak, Y. S., Um, S. Y., Son, T. G., & Kim, D. J. (2008). Effect of regular exercise on senile dementia patients. *International Journal of Sports Medicine, 29*(06), 471–474. https://doi.org/10.1055/s-2007-964853

Lam, L. C. W., Chan, W. C., Leung, T., Fung, A. W. T., & Leung, E. M. F. (2015). Would older adults with mild cognitive impairment adhere to and benefit from a structured lifestyle activity intervention to enhance cognition?: a cluster randomized controlled trial. *PloS ONE, 10*(3), e0118173. https://doi.org/10.1371/journal.pone.0118173

Lam, L. C., Chau, R. C., Wong, B. M., Fung, A. W., Tam, C. W., Leung, G. T., ... & Chan, W. M. (2012). A 1-year randomized controlled trial comparing mind body exercise (Tai Chi) with stretching and toning exercise on cognitive function in older Chinese adults at risk of cognitive decline. *Journal of the American Medical Directors Association, 13*(6), 568-e15. https://doi.org/10.1016/j.jamda.2012.03.008

Langlois, F., Vu, T. T. M., Chassé, K., Dupuis, G., Kergoat, M. J., & Bherer, L. (2012). Benefits of physical exercise training on cognition and quality of life in frail older adults. *Journals of Gerontology Series B: Psychological Sciences and Social Sciences, 68*(3), 400–404. https://doi.org/10.1093/geronb/gbs069

Lautenschlager, N. T., Cox, K. L., Flicker, L., Foster, J. K., Van Bockxmeer, F. M., Xiao, J., ... & Almeida, O. P. (2008). Effect of physical activity on cognitive function in older adults at risk for Alzheimer disease: a randomized trial. *JAMA, 300*(9), 1027–1037. https://doi.org/10.1001/jama.300.9.1027

Lazarou, I., Parastatidis, T., Tsolaki, A., Gkioka, M., Karakostas, A., Douka, S., & Tsolaki, M. (2017). International ballroom dancing against neurodegeneration: a randomized controlled trial in Greek community-dwelling elders with mild cognitive impairment. *American Journal of Alzheimer's Disease & Other Dementias, 32*(8), 489–499. https://doi.org/10.1177/1533317517725813

Liu, J. H., Alderman, B. L., Song, T. F., Chen, F. T., Hung, T. M., & Chang, Y. K. (2018). A randomized controlled trial of coordination exercise on cognitive function in obese adolescents. *Psychology of Sport and Exercise, 34*, 29–38. https://doi.org/10.1016/j.psychsport.2017.09.003

Liu-Ambrose, T., Best, J. R., Davis, J. C., Eng, J. J., Lee, P. E., Jacova, C., ... & Hsiung, G. Y. R. (2016). Aerobic exercise and vascular cognitive impairment: a randomized controlled trial. *Neurology, 87*(20), 2082–2090. https://doi.org/10.1212/WNL.0000000000003332

Lord, S. R., Castell, S., Corcoran, J., Dayhew, J., Matters, B., Shan, A., & Williams, P. (2003). The effect of group exercise on physical functioning and falls in frail older people living in retirement villages: a randomized, controlled trial. *Journal of the American Geriatrics Society, 51*(12), 1685–1692. https://doi.org/10.1046/j.1532-5415.2003.51551.x

Lü, J., Sun, M., Liang, L., Feng, Y., Pan, X., & Liu, Y. (2016). Effects of momentum-based dumbbell training on cognitive function in older adults with mild cognitive impairment: a pilot randomized controlled trial. *Clinical Interventions in Aging, 11*, 9–16. https://doi.org/10.2147/cia.s96042

Marmeleira, J. F., Godinho, M. B., & Fernandes, O. M. (2009). The effects of an exercise program on several abilities associated with driving performance in older adults. *Accident Analysis & Prevention, 41*(1), 90–97. https://doi.org/10.1016/j.aap.2008.09.008

Matsufuji, S., Shoji, T., Yano, Y., Tsujimoto, Y., Kishimoto, H., Tabata, T., ... & Inaba, M. (2015). Effect of chair stand exercise on activity of daily living: a randomized controlled trial in hemodialysis patients. *Journal of Renal Nutrition, 25*(1), 17–24. https://doi.org/10.1053/j.jrn.2014.06.010

Mavros, Y., Gates, N., Wilson, G. C., Jain, N., Meiklejohn, J., Brodaty, H., ... & Fiatarone Singh, M. A. (2017). Mediation of cognitive function improvements by strength gains after resistance training in older adults with mild cognitive impairment: outcomes of the study of mental and resistance training. *Journal of the American Geriatrics Society, 65*(3), 550–559. https://doi.org/10.1111/jgs.14542

McCann, I. L., & Holmes, D. S. (1984). Influence of aerobic exercise on depression. *Journal of Personality and Social Psychology, 46*(5), 1142–1147. https://doi.org/10.1037/0022-3514.46.5.1142

McNeil, J. K., LeBlanc, E. M., & Joyner, M. (1991). The effect of exercise on depressive symptoms in the moderately depressed elderly. *Psychology and Aging, 6*(3), 487–488. https://doi.org/10.1037/0882-7974.6.3.487

Merom, D., Mathieu, E., Cerin, E., Morton, R. L., Simpson, J. M., Rissel, C., ... & Cumming, R. G. (2016). Social dancing and incidence of falls in older adults: a cluster randomised controlled trial. *PLoS Medicine, 13*(8), e1002112. https://doi.org/10.1371/journal.pmed.1002112

Middleton, L. E., Ventura, M. I., Santos-Modesitt, W., Poelke, G., Yaffe, K., & Barnes, D. E. (2018). The Mental Activity and eXercise (MAX) trial: Effects on physical function and quality of life among older adults with cognitive complaints. *Contemporary Clinical Trials, 64*, 161–166. https://doi.org/10.1016/j.cct.2017.10.009

Morris, J. K., Vidoni, E. D., Johnson, D. K., Van Sciver, A., Mahnken, J. D., Honea, R. A., ... & Burns, J. M. (2017). Aerobic exercise for Alzheimer's disease: A randomized controlled pilot trial. *PloS ONE, 12*(2), e0170547. https://doi.org/10.1371/journal.pone.0170547

Munguía-Izquierdo, D., & Legaz-Arrese, A. (2008). Assessment of the effects of aquatic therapy on global symptomatology in patients with fibromyalgia syndrome: a randomized controlled trial. *Archives of Physical Medicine and Rehabilitation, 89*(12), 2250–2257. https://doi.org/10.1016/j.apmr.2008.03.026

Nagamatsu, L. S., Chan, A., Davis, J. C., Beattie, B. L., Graf, P., Voss, M. W., ... & Liu-Ambrose, T. (2013). Physical activity improves verbal and spatial memory in older adults with probable mild cognitive impairment: a 6-month randomized controlled trial. *Journal of Aging Research, 2013*. https://doi.org/10.1155/2013/861893

Nagamatsu, L. S., Handy, T. C., Hsu, C. L., Voss, M., & Liu-Ambrose, T. (2012). Resistance training promotes cognitive and functional brain plasticity in seniors with probable mild cognitive impairment. *Archives of Internal Medicine, 172*(8), 666–668. https://doi.org/10.1001/archinternmed.2012.379

Napoli, N., Shah, K., Waters, D. L., Sinacore, D. R., Qualls, C., & Villareal, D. T. (2014). Effect of weight loss, exercise, or both on cognition and quality of life in obese older adults. *The American Journal of Clinical Nutrition, 100*(1), 189–198. https://doi.org/10.3945/ajcn.113.082883

Nascimento, C. M. C., Pereira, J. R., Pires de Andrade, L., Garuffi, M., Ayan, C., Kerr, D. S., ... & Stella, F. (2015). Physical exercise improves peripheral BDNF levels and cognitive functions in mild cognitive impairment elderly with different bdnf Val66Met genotypes. *Journal of Alzheimer's Disease, 43*(1), 81–91. https://doi.org/10.3233/JAD-140576

Nascimento, C. M. C., Rodrigues Pereira, J., Pires de Andrade, L., Garuffi, M., Leme Talib, L., Vicente Forlenza, O., ... & Stella, F. (2014). Physical exercise in MCI elderly promotes reduction of pro-inflammatory cytokines and improvements on cognition and BDNF peripheral levels. *Current Alzheimer Research, 11*(8), 799–805. http://dx.doi.org/10.2174/1567205011081409101228490

Oken, B. S., Kishiyama, S., Zajdel, D., Bourdette, D., Carlsen, J., Haas, M., ... & Mass, M. (2004). Randomized controlled trial of yoga and exercise in multiple sclerosis. *Neurology, 62*(11), 2058–2064. https://doi.org/10.1212/01.WNL.0000129534.88602.5C

Pahor, M., Blair, S. N., Espeland, M., Fielding, R., Gill, T. M., Guralnik, J. M., Maraldi, C., Miller, M. E., Newman, A. B., & Rejeski, W. J., (2006). Effects of a physical activity intervention on measures of physical performance: results of the lifestyle interventions and independence for elders pilot (LIFE-P) study. *Journals of Gerontology Series A: Biological Sciences and Medical Sciences, 61*(11), 1157–1165. https://doi.org/10.1093/gerona/61.11.1157

Pahor, M., Guralnik, J. M., Ambrosius, W. T., Blair, S., Bonds, D. E., Church, T. S., ... & LIFE Study Investigators. (2014). Effect of structured physical activity on prevention of major mobility disability in older adults: the LIFE study randomized clinical trial. *JAMA, 311*(23), 2387–2396. https://doi.org/10.1001/jama.2014.5616

Pierce, T. W., Madden, D. J., Siegel, W. C., & Blumenthal, J. A. (1993). Effects of aerobic exercise on cognitive and psychosocial functioning in patients with mild hypertension. *Health Psychology, 12*(4), 286–291. https://doi.org/10.1037/0278-6133.12.4.286

Poinsatte, K., Smith, E. E., Torres, V. O., Ortega, S. B., Huebinger, R. M., Cullum, C. M., ... & Stowe, A. M. (2019). T and B cell subsets differentially correlate with amyloid deposition and neurocognitive function in patients with amnestic mild cognitive impairment after one year of physical activity. *Exercise Immunology Review, 25*, 34–49.

Prehn, K., Lesemann, A., Krey, G., Witte, A. V., Köbe, T., Grittner, U., & Flöel, A. (2019). Using resting-state fMRI to assess the effect of aerobic exercise on functional connectivity of the DLPFC in older overweight adults. *Brain and Cognition, 131*, 34–44. https://doi.org/10.1016/j.bandc.2017.08.006

Roth, D. L., & Holmes, D. S. (1987). Influence of aerobic exercise training and relaxation training on physical and psychologic health following stressful life events. *Psychosomatic Medicine, 49*(4), 355–365. https://doi.org/10.1097/00006842-198707000-00004

Ruiz, J. R., Gil-Bea, F., Bustamante-Ara, N., Rodríguez-Romo, G., Fiuza-Luces, C., Serra-Rexach, J. A., ... & Lucia, A. (2015). Resistance training does not have an effect on cognition or related serum biomarkers in nonagenarians: a randomized controlled trial. *International Journal of Sports Medicine, 36*(01), 54–60. https://doi.org/10.1055/s-0034-1375693

Scherder, E. J., Van Paasschen, J., Deijen, J. B., Van Der Knokke, S., Orlebeke, J. F. K., Burgers, I., ... & Sergeant, J. A. (2005). Physical activity and executive functions in the elderly with mild cognitive impairment. *Aging & Mental Health, 9*(3), 272–280. https://doi.org/10.1080/13607860500089930

Schmidt, M., Jäger, K., Egger, F., Roebers, C. M., & Conzelmann, A. (2015). Cognitively engaging chronic physical activity, but not aerobic exercise, affects executive functions in primary school children: a group-randomized controlled trial. *Journal of Sport and Exercise Psychology, 37*(6), 575–591. https://doi.org/10.1123/jsep.2015-0069

Stuckenschneider, T., Sanders, M. L., Devenney, K. E., Aaronson, J. A., Abeln, V., Claassen, J. A., ... & Schneider, S. (2021). NeuroExercise: the effect of a 12-month exercise intervention on cognition in mild cognitive impairment—a multicenter randomized controlled trial. *Frontiers in Aging Neuroscience, 12*, 621947. https://doi.org/10.3389/fnagi.2020.621947

Sugano, K., Yokogawa, M., Yuki, S., Dohmoto, C., Yoshita, M., Hamaguchi, T., ... & Yamada, M. (2012). Effect of cognitive and aerobic training intervention on older adults with mild or no cognitive impairment: a derivative study of the nakajima project. *Dementia and Geriatric Cognitive Disorders Extra, 2*(1), 69–80. https://doi.org/10.1159/000337224

Sungkarat, S., Boripuntakul, S., Kumfu, S., Lord, S. R., & Chattipakorn, N. (2018). Tai Chi improves cognition and plasma BDNF in older adults with mild cognitive impairment: a randomized controlled trial. *Neurorehabilitation and Neural Repair, 32*(2), 142–149. https://doi.org/10.1177/1545968317753682

Suzuki, T., Shimada, H., Makizako, H., Yoshida, D., Tsutsumimoto, K., Anan, Y., ... & Park, H. (2012). Effects of multicomponent exercise on cognitive function in older adults with amnestic mild cognitive impairment: a randomized controlled trial. *BMC Neurology, 12*(1), 1–9. https://doi.org/10.1186/1471-2377-12-128

Tarazona-Santabalbina, F. J., Gómez-Cabrera, M. C., Pérez-Ros, P., Martínez-Arnau, F. M., Cabo, H., Tsaparas, K., ... & Viña, J. (2016). A multicomponent exercise intervention that reverses frailty and improves cognition, emotion, and social networking in the community-dwelling frail elderly: a randomized clinical trial. *Journal of the American Medical Directors Association, 17*(5), 426–433. https://doi.org/10.1016/j.jamda.2016.01.019

Tarumi, T., Rossetti, H., Thomas, B. P., Harris, T., Tseng, B. Y., Turner, M., ... & Zhang, R. (2019). Exercise training in amnestic mild cognitive impairment: a one-year randomized controlled trial. *Journal of Alzheimer's Disease, 71*(2), 421–433. doi: 10.3233/JAD-181175

Telenius, E. W., Engedal, K., & Bergland, A. (2015). Long-term effects of a 12 weeks high-intensity functional exercise program on physical function and mental health in nursing home residents with dementia: a single blinded randomized controlled trial. *BMC Geriatrics, 15*(1), 1–11. https://doi.org/10.1186/s12877-015-0151-8

Ten Brinke, L. F., Bolandzadeh, N., Nagamatsu, L. S., Hsu, C. L., Davis, J. C., Miran-Khan, K., & Liu-Ambrose, T. (2015). Aerobic exercise increases hippocampal volume in older women with probable mild cognitive impairment: a 6-month randomised controlled trial. *British Journal of Sports Medicine, 49*(4), 248–254. https://doi.org/10.1136/bjsports- 2013-093184

Tench, C. M., McCarthy, J., McCurdie, I., White, P. D., & D'Cruz, D. P. (2003). Fatigue in systemic lupus erythematosus: a randomized controlled trial of exercise. *Rheumatology, 42*(9), 1050–1054. https://doi.org/10.1093/rheumatology/keg289

Thomas, B. P., Tarumi, T., Sheng, M., Tseng, B., Womack, K. B., Cullum, C. M., ... & Lu, H. (2020). Brain perfusion change in patients with mild cognitive impairment after 12 months of aerobic exercise training. *Journal of Alzheimer's Disease, 75*(2), 617–631. doi: 10.3233/JAD-190977

Tsai, P. F., Chang, J. Y., Beck, C., Kuo, Y. F., & Keefe, F. J. (2013). A pilot cluster-randomized trial of a 20-week Tai Chi program in elders with cognitive impairment and osteoarthritic knee: effects on pain and other health outcomes. *Journal of Pain and Symptom Management, 45*(4), 660–669. https://doi.org/10.1016/j.jpainsymman.2012.04.009

van De Rest, O., van der Zwaluw, N. L., Tieland, M., Adam, J. J., Hiddink, G. J., Van Loon, L. J., & de Groot, L. C. (2014). Effect of resistance-type exercise training with or without protein supplementation on cognitive functioning in frail and pre-frail elderly: secondary analysis of a randomized, double-blind, placebo-controlled trial. *Mechanisms of Ageing and Development, 136*, 85–93. https://doi.org/10.1016/j.mad.2013.12.005

van Uffelen, J. G., Chinapaw, M. J., van Mechelen, W., & Hopman-Rock, M. (2008). Walking or vitamin B for cognition in older adults with mild cognitive impairment? A randomised controlled trial. *British Journal of Sports Medicine, 42* (5), 344–351. https://doi.org/10.1136/bjsm.2007.044735

Varela, S., Ayán, C., Cancela, J. M., & Martín, V. (2012). Effects of two different intensities of aerobic exercise on elderly people with mild cognitive impairment: a randomized pilot study. *Clinical Rehabilitation, 26*(5), 442–450. https://doi.org/10.1177/0269215511425835

Venturelli, M., Scarsini, R., & Schena, F. (2011). Six-month walking program changes cognitive and ADL performance in patients with Alzheimer. *American Journal of Alzheimer's Disease & Other Dementias, 26*(5), 381–388. https://doi.org/10.1177/1533317511418956

Villareal, D. T., Chode, S., Parimi, N., Sinacore, D. R., Hilton, T., Armamento-Villareal, R., ... & Shah, K. (2011). Weight loss, exercise, or both and physical function in obese older adults. *New England Journal of Medicine, 364*(13), 1218–1229. https://doi.org/10.1056/NEJMoa1008234

Wallman, K. E., Morton, A. R., Goodman, C., Grove, R., & Guilfoyle, A. M. (2004). Randomised controlled trial of graded exercise in chronic fatigue syndrome. *Medical Journal of Australia, 180*(9), 444–448. https://doi.org/10.5694/j.1326-5377.2004.tb06019.x

Wei, X. H., & Ji, L. L. (2014). Effect of handball training on cognitive ability in elderly with mild cognitive impairment. *Neuroscience Letters, 566*, 98–101. https://doi.org/10.1016/j.neulet.2014.02.035

Williamson, J. D., Espeland, M., Kritchevsky, S. B., Newman, A. B., King, A. C., Pahor, M., ... & LIFE Study Investigators. (2009). Changes in cognitive function in a randomized trial of physical activity: results of the lifestyle interventions and independence for elders pilot study. *Journals of Gerontology Series A: Biomedical Sciences and Medical Sciences, 64*(6), 688–694. https://doi.org/10.1093/gerona/glp014

Yogev-Seligmann, G., Eisenstein, T., Ash, E., Giladi, N., Sharon, H., Nachman, S., ... & Lerner, Y. (2021). Neurocognitive plasticity is associated with cardiorespiratory fitness following physical exercise in older adults with amnestic mild cognitive impairment. *Journal of Alzheimer's Disease, 81*(1), 91–112. doi: 10.3233/JAD-201429

Yoon, D. H., Kang, D., Kim, H. J., Kim, J. S., Song, H. S., & Song, W. (2017). Effect of elastic band-based high-speed power training on cognitive function, physical performance and muscle strength in older women with mild cognitive impairment. *Geriatrics & Gerontology International, 17*(5), 765–772. https://doi.org/10.1111/ggi.12784

Zhu, Y., Wu, H., Qi, M., Wang, S., Zhang, Q., Zhou, L., ... & Wang, T. (2018). Effects of a specially designed aerobic dance routine on mild cognitive impairment. Clinical Interventions in Aging, 13, 1691–1700. https://doi.org/10.2147/CIA.S163067

**B. Using a Different Type of Physical Intervention**


Aadland, K. N., Ommundsen, Y., Anderssen, S. A., Brønnick, K. S., Moe, V. F., Resaland, G. K., ... & Aadland, E. (2019). Effects of the Active Smarter Kids (ASK) physical activity school-based intervention on executive functions: a cluster-randomized controlled trial. Scandinavian *Journal of Educational Research, 63*(2), 214–228. https://doi.org/10.1080/00313831.2017.1336477

Bantoft, C., Summers, M. J., Tranent, P. J., Palmer, M. A., Cooley, P. D., & Pedersen, S. J. (2016). Effect of standing or walking at a workstation on cognitive function: a randomized counterbalanced trial. *Human factors, 58*(1), 140–149. https://doi.org/10.1177/0018720815605446

Butzer, B., Van Over, M., Noggle Taylor, J. J., & Khalsa, S. B. S. (2015). Yoga may mitigate decreases in high school grades. *Evidence-Based Complementary and Alternative Medicine, 2015*. https://doi.org/10.1155/2015/259814

Chang, Y. K., Tsai, Y. J., Chen, T. T., & Hung, T. M. (2013). The impacts of coordinative exercise on executive function in kindergarten children: an ERP study. *Experimental Brain Research, 225*(2), 187–196. https://doi.org/10.1007/s00221-012-3360-9

Duarte, L., Gonçalves, M., Mendes, P., Matos, L. C., Greten, H. J., & Machado, J. (2020). Can Qigong improve attention in adolescents? A prospective randomised controlled trial. *Journal of Bodywork and Movement Therapies, 24*(1), 175–181. https://doi.org/10.1016/j.jbmt.2019.05.005

Eggenberger, P., Wolf, M., Schumann, M., & de Bruin, E. D. (2016). Exergame and balance training modulate prefrontal brain activity during walking and enhance executive function in older adults. *Frontiers in Aging Neuroscience, 8*, 66. https://doi.org/10.3389/fnagi.2016.00066

Frändin, K., Grönstedt, H., Helbostad, J. L., Bergland, A., Andresen, M., Puggaard, L., ... & Hellström, K. (2016). Long-term effects of individually tailored physical training and activity on physical function, well-being and cognition in Scandinavian nursing home residents: a randomized controlled trial. *Gerontology, 62*(6), 571–580. https://doi.org/10.1159/000443611

Gothe, N. P., Keswani, R. K., & McAuley, E. (2016). Yoga practice improves executive function by attenuating stress levels. *Biological Psychology, 121*(Pt A), 109–116. https://doi.org/10.1016/j.biopsycho.2016.10.010.

Gothe, N. P., Kramer, A. F., & McAuley, E. (2017). Hatha yoga practice improves attention and processing speed in older adults: results from an 8-week randomized control trial. *The Journal of Alternative and Complementary Medicine, 23*(1), 35–40. https://doi.org/10.1089/acm.2016.0185

Hagins, M., & Rundle, A. (2016). Yoga improves academic performance in urban high school students compared to physical education: a randomized controlled trial. *Mind, Brain, and Education, 10*(2), 105–116. https://doi.org/10.1111/mbe.12107

Have, M., Nielsen, J. H., Ernst, M. T., Gejl, A. K., Fredens, K., Grøntved, A., & Kristensen, P. L. (2018). Classroom-based physical activity improves children's math achievement–A randomized controlled trial. *PloS ONE, 13*(12), e0208787. https://doi.org/10.1371/journal.pone.0208787

Hong, S. G., Kim, J. H., & Jun, T. W. (2018). Effects of 12-week resistance exercise on electroencephalogram patterns and cognitive function in the elderly with mild cognitive impairment: a randomized controlled trial. *Clinical Journal of Sport Medicine, 28*(6), 500–508. doi: 10.1097/JSM.0000000000000476

Kalbe, E., Roheger, M., Paluszak, K., Meyer, J., Becker, J., Fink, G. R., ... & Kessler, J. (2018). Effects of a cognitive training with and without additional physical activity in healthy older adults: a follow-up 1 year after a randomized controlled trial. Frontiers in Aging Neuroscience, 10, 407. https://doi.org/10.3389/fnagi.2018.00407

Kattenstroth, J. C., Kalisch, T., Holt, S., Tegenthoff, M., & Dinse, H. R. (2013). Six months of dance intervention enhances postural, sensorimotor, and cognitive performance in elderly without affecting cardio-respiratory functions. *Frontiers in Aging Neuroscience, 5*, 5. https://doi.org/10.3389/fnagi.2013.00005

Kauts, A., & Sharma, N. (2009). Effect of yoga on academic performance in relation to stress. *International Journal of Yoga, 2*(1), 39–43. https://doi.org/10.4103/0973-6131.53860

Lakes, K. D., & Hoyt, W. T. (2004). Promoting self-regulation through school-based martial arts training. *Journal of Applied Developmental Psychology, 25*(3), 283–302. https://doi.org/10.1016/j.appdev.2004.04.002

Maillot, P., Perrot, A., & Hartley, A. (2012). Effects of interactive physical-activity video-game training on physical and cognitive function in older adults. *Psychology and Aging, 27*(3), 589–600. https://doi.org/10.1037/a0026268

Matson, T. E., Anderson, M. L., Renz, A. D., Greenwood-Hickman, M. A., McClure, J. B., & Rosenberg, D. E. (2019). Changes in self-reported health and psychosocial outcomes in older adults enrolled in sedentary behavior intervention study. *American Journal of Health Promotion, 33*(7), 1053–1057. https://doi.org/10.1177/0890117119841405

Mavilidi, M. F., Lubans, D. R., Eather, N., Morgan, P. J., & Riley, N. (2018). Preliminary efficacy and feasibility of “Thinking While Moving in English”: A program with physical activity integrated into primary school English lessons. *Children, 5*(8), 109. https://doi.org/10.3390/children5080109

Nguyen, M. H., & Kruse, A. (2012). A randomized controlled trial of Tai chi for balance, sleep quality and cognitive performance in elderly Vietnamese. *Clinical Interventions in Aging, 7*, 185–190. https://doi.org/10.2147/CIA.S32600

Norouzi, E., Vaezmosavi, M., Gerber, M., Pühse, U., & Brand, S. (2019). Dual-task training on cognition and resistance training improved both balance and working memory in older people. *The Physician and Sports Medicine, 47*(4), 471–478. https://doi.org/10.1080/00913847.2019.1623996

Nishiguchi, S., Yamada, M., Tanigawa, T., Sekiyama, K., Kawagoe, T., Suzuki, M., ... & Tsuboyama, T. (2015). A 12-week physical and cognitive exercise program can improve cognitive function and neural efficiency in community-dwelling older adults: a randomized controlled trial. *Journal of the American Geriatrics Society, 63*(7), 1355–1363. https://doi.org/10.1111/jgs.13481

Palleschi, L., Vetta, F., De Gennaro, E., Idone, G., Sottosanti, G., Gianni, W., & Marigliano, V. (1996). Effect of aerobic training on the cognitive performance of elderly patients with senile dementia of Alzheimer type. *Archives of Gerontology and Geriatrics, 22*, 47–50. https://doi.org/10.1016/0167-4943(96)86912-3

Pesce, C., Masci, I., Marchetti, R., Vazou, S., Sääkslahti, A., & Tomporowski, P. D. (2016). Deliberate play and preparation jointly benefit motor and cognitive development: mediated and moderated effects. *Frontiers in Psychology, 7*, 349. https://doi.org/10.3389/fpsyg.2016.00349

Pichierri, G., Coppe, A., Lorenzetti, S., Murer, K., & de Bruin, E. D. (2012). The effect of a cognitive-motor intervention on voluntary step execution under single and dual task conditions in older adults: a randomized controlled pilot study. *Clinical Interventions in Aging, 7*, 175–184. http://dx.doi.org/10.2147/CIA.S32558

Powell, R. R. (1974). Psychological effects of exercise therapy upon institutionalized geriatric mental patients. *Journal of Gerontology, 29*(2), 157–161. https://doi.org/10.1093/geronj/29.2.157

Purohit, S. P., & Pradhan, B. (2017). Effect of yoga program on executive functions of adolescents dwelling in an orphan home: A randomized controlled study. *Journal of Traditional and Complementary Medicine, 7*(1), 99–105. https://doi.org/10.1016/j.jtcme.2016.03.001

Schättin, A., Arner, R., Gennaro, F., & de Bruin, E. D. (2016). Adaptations of prefrontal brain activity, executive functions, and gait in healthy elderly following exergame and balance training: a randomized-controlled study. *Frontiers in Aging Neuroscience, 8*, 278. https://doi.org/10.3389/fnagi.2016.00278

Schoene, D., Lord, S. R., Delbaere, K., Severino, C., Davies, T. A., & Smith, S. T. (2013). A randomized controlled pilot study of home-based step training in older people using videogame technology. *PloS ONE, 8*(3), e57734. https://doi.org/10.1371/journal.pone.0057734

Schoene, D., Valenzuela, T., Toson, B., Delbaere, K., Severino, C., Garcia, J., ... & Lord, S. R. (2015). Interactive cognitive-motor step training improves cognitive risk factors of falling in older adults–a randomized controlled trial. *PLoS ONE, 10*(12), e0145161. https://doi.org/10.1371/journal.pone.0145161

Sharma, V. K., Subramanian, S. K., Arunachalam, V., Radhakrishnan, K., Ramamurthy, S., & Ravindran, B. S. (2017). Auditory and visual reaction times in school going adolescents: Effect of structured and unstructured physical training–a randomized control trial. *International Journal of Adolescent Medicine and Health, 29*(4). https://doi.org/10.1515/ijamh-2015-0060

Tao, J., Liu, J., Egorova, N., Chen, X., Sun, S., Xue, X., ... & Kong, J. (2016). Increased hippocampus–medial prefrontal cortex resting-state functional connectivity and memory function after Tai Chi Chuan practice in elder adults. *Frontiers in Aging Neuroscience, 8*, 25. https://doi.org/10.3389/fnagi.2016.00025

van den Berg, V., Singh, A. S., Komen, A., Hazelebach, C., van Hilvoorde, I., & Chinapaw, M. J. (2019). Integrating juggling with math lessons: A randomized controlled trial assessing effects of physically active learning on maths performance and enjoyment in primary school children. *International Journal of Environmental Research and Public Health, 16*(14), 2452. https://doi.org/10.3390/ijerph16142452

### C. Non-published study

Li, S. Z. (2016). Effects of Baduanjin on global cognitive function and memory in the elderly with mild cognitive impairment. Thesis Dissertation.

Russell, E. M. (1983). The effects of an aerobic conditioning program on reaction times of older sedentary adults. Thesis Dissertation.

Shan, Y. T. (2016). The effects of “24-style” simplified Taijiquan on memory, attention and executive function of the middle-aged and elderly in community—a randomized controlled study. Thesis Dissertation.

Yang, Y. (2019). Effects of 8-week Taijiquan on cognitive control and working memory of the elderly in community. Thesis Dissertation.

**D. No Cognitive Outcome**


Niemann, C., Godde, B., & Voelcker-Rehage, C. (2014). Not only cardiovascular, but also coordinative exercise increases hippocampal volume in older adults. *Frontiers in Aging Neuroscience, 6*, 170. https://doi.org/10.3389/fnagi.2014.00170

Zheng, G., Lan, X., Li, M., Ling, K., Lin, H., Chen, L., ... & Fang, Q. (2015). Effectiveness of Tai Chi on physical and psychological health of college students: Results of a randomized controlled trial. *PloS ONE, 10*(7), e0132605. https://doi.org/10.1371/journal.pone.0132605

**E. Not a Randomized Controlled Trial**


Barry, A. J., Steinmetz, J. R., Page, H. F., & Rodahl, K. (1966). The effects of physical conditioning on older individuals. II. Motor performance and cognitive function. Journal of Gerontology, 21(2), 192–199. https://doi.org/10.1093/geronj/21.2.192

Dustman, R. E., Ruhling, R. O., Russell, E. M., Shearer, D. E., Bonekat, H. W., Shigeoka, J. W., ... & Bradford, D. C. (1984). Aerobic exercise training and improved neuropsychological function of older individuals. *Neurobiology of Aging, 5*(1), 35–42. https://doi.org/10.1016/0197-4580(84)90083-6

Hassmén, P., Ceci, R., & Bäckman, L. (1992). Exercise for older women: a training method and its influences on physical and cognitive performance. *European Journal of Applied Physiology and Occupational Physiology, 64*(5), 460–466. https://doi.org/10.1007/BF00625068

Hassmén, P., & Koivula, N. (1997). Mood, physical working capacity and cognitive performance in the elderly as related to physical activity. *Aging Clinical and Experimental Research, 9*(1), 136–142. https://doi.org/10.1007/BF03340139

Hill, R. D., Storandt, M., & Malley, M. (1993). The impact of long-term exercise training on psychological function in older adults. *Journal of Gerontology, 48*(1), P12–P17. https://doi.org/10.1093/geronj/48.1.P12

Perri, S., & Templer, D. I. (1985). The effects of an aerobic exercise program on psychological variables in older adults. *The International Journal of Aging and Human Development, 20*(3), 167–172. https://doi.org/10.2190/A6AB-MBN1-G8HV-258H

Predovan, D., Fraser, S. A., Renaud, M., & Bherer, L. (2012). The effect of three months of aerobic training on stroop performance in older adults. *Journal of Aging Research*, 2012. https://doi.org/10.1155/2012/269815

Rikli, R. E., & Edwards, D. J. (1991). Effects of a three-year exercise program on motor function and cognitive processing speed in older women. *Research Quarterly for Exercise and Sport, 62*(1), 61–67. https://doi.org/10.1080/02701367.1991.10607519

Sjöwall, D., Thorell, L. B., Mandic, M., & Westerståhl, M. (2019). No effects of a long-term physical activity intervention on executive functioning among adolescents. *SAGE Open Medicine, 7*, 2050312119880734. https://doi.org/10.1177/2050312119880734

Smiley-Oyen, A. L., Lowry, K. A., Francois, S. J., Kohut, M. L., & Ekkekakis, P. (2008). Exercise, fitness, and neurocognitive function in older adults: the “selective improvement” and “cardiovascular fitness” hypotheses. *Annals of Behavioral Medicine, 36*(3), 280–291. https://doi.org/10.1007/s12160-008-9064-5

Spitzer, U. S., & Hollmann, W. (2013). Experimental observations of the effects of physical exercise on attention, academic and prosocial performance in school settings. *Trends in Neuroscience and Education, 2*(1), 1–6. https://doi.org/10.1016/j.tine.2013.03.002

Stroth, S., Reinhardt, R. K., Thöne, J., Hille, K., Schneider, M., Härtel, S., ... & Spitzer, M. (2010). Impact of aerobic exercise training on cognitive functions and affect associated to the COMT polymorphism in young adults. *Neurobiology of Learning and Memory, 94*(3), 364–372. https://doi.org/10.1016/j.nlm.2010.08.003

Venckunas, T., Snieckus, A., Trinkunas, E., Baranauskiene, N., Solianik, R., Juodsnukis, A., ... & Kamandulis, S. (2016). Interval running training improves cognitive flexibility and aerobic power of young healthy adults. *Journal of Strength and Conditioning Research, 30*(8), 2114–2121. https://doi.org/10.1519/JSC.0000000000001322

Woost, L., Bazin, P. L., Taubert, M., Trampel, R., Tardif, C. L., Garthe, A., ... & Klein, T. A. (2018). Physical exercise and spatial training: a longitudinal study of effects on cognition, growth factors, and hippocampal plasticity. *Scientific Reports, 8*(1), 1–13. https://doi.org/10.1038/s41598-018-19993-9

### F. Less Than One Week of Training


Basso, J. C., Shang, A., Elman, M., Karmouta, R., & Suzuki, W. A. (2015). Acute exercise improves prefrontal cortex but not hippocampal function in healthy adults. *Journal of the International Neuropsychological Society, 21*(10), 791–801. https://doi.org/10.1017/S135561771500106X

Budde, H., Voelcker-Rehage, C., Pietrassyk-Kendziorra, S., Machado, S., Ribeiro, P., & Arafat, A. M. (2010). Steroid hormones in the saliva of adolescents after different exercise intensities and their influence on working memory in a school setting. *Psychoneuroendocrinology, 35*(3), 382–391. https://doi.org/10.1016/j.psyneuen.2009.07.015

Hogan, C. L., Mata, J., & Carstensen, L. L. (2013). Exercise holds immediate benefits for affect and cognition in younger and older adults. *Psychology and Aging, 28*(2), 587–594. https://doi.org/10.1037/a0032634

Howie, E. K., Beets, M. W., & Pate, R. R. (2014). Acute classroom exercise breaks improve on-task behavior in 4th and 5th grade students: a dose–response. *Mental Health and Physical Activity, 7*(2), 65–71. https://doi.org/10.1016/j.mhpa.2014.05.002

Janssen, M., Chinapaw, M. J. M., Rauh, S. P., Toussaint, H. M., Van Mechelen, W., & Verhagen, E. A. L. M. (2014). A short physical activity break from cognitive tasks increases selective attention in primary school children aged 10–11. *Mental Health and Physical Activity, 7*(3), 129–134. https://doi.org/10.1016/j.mhpa.2014.07.001

Pinto-Escalona, T., & Martínez-de-Quel, Ó. (2019). Ten minutes of interdisciplinary physical activity improve academic performance. *Apunts. Educació Física i Esports, 138*(4), 82–94. https://dx.doi.org/10.5672/apunts.2014-0983.cat.(2019/4).138.07

**G. Information not available**

Baniqued, P. L., Gallen, C. L., Voss, M. W., Burzynska, A. Z., Wong, C. N., Cooke, G. E., ... & D'Esposito, M. (2018). Brain network modularity predicts exercise-related executive function gains in older adults. *Frontiers in Aging Neuroscience, 9*, 426. https://doi.org/10.3389/fnagi.2017.00426

**H. Lack of control group**

Chapman, S. B., Aslan, S., Spence, J. S., Keebler, M. W., DeFina, L. F., Didehbani, N., Perez, A. M., Lu, H., & D'Esposito, M. (2016). Distinct Brain and Behavioral Benefits from Cognitive vs. Physical Training: A Randomized Trial in Aging Adults. *Frontiers in Human Neuroscience, 10*, 338. https://doi.org/10.3389/fnhum.2016.00338

Lennemann, L. M., Sidrow, K. M., Johnson, E. M., Harrison, C. R., Vojta, C. N., & Walker, T. B. (2013). The influence of agility training on physiological and cognitive performance. *The Journal of Strength & Conditioning Research, 27*(12), 3300–3309. doi: 10.1519/JSC.0b013e31828ddf06

**I. Only posttest cognitive assessment**

Lakes, K. D., Bryars, T., Sirisinahal, S., Salim, N., Arastoo, S., Emmerson, N., ... & Kang, C. J. (2013). The healthy for life taekwondo pilot study: a preliminary evaluation of effects on executive function and BMI, feasibility, and acceptability. *Mental Health and Physical Activity, 6*(3), 181–188. https://doi.org/10.1016/j.mhpa.2013.07.002

**Table S1.** Characteristics of the meta-analyses included in the umbrella review.

| Meta-analysis | Type of exercise | Age range | Cognitive domains | Other inclusion criteria | Studies originally included | Studies included in the umbrella review | Effect sizes included in the umbrella review | Original final effect | Estimated final effect | Heterogeneity ($I^2$) | Average participants per effect size | Average program duration (months) | Dependence dealing |
|---|---|---|---|---|---|---|---|---|---|---|---|---|---|
| Aghjayan et al. (2022) | Aerobic | Older<br>(Mean ≥ 55 years) | Episodic memory | | 36 | 17 | 45 | 0.28 | 0.20 [0.07, 0.34] | 44.89 | 54<br>(range 15–133) | 6.3<br>(range 2–24) | None / not reported |
| Amatriain-Fernández et al. (2021) | Multiple | Children & adolescents<br>(1–19 years) | Executive control | • Only cognitively and physically healthy<br>• Program duration ≥ 6 weeks<br>• Session duration ≥ 10 min | 10 | 8 | 18 | 0.06 | 0.09 [−0.01, 0.19] | 47.63 | 271<br>(range 23–584) | 7.2<br>(range 1.5–24) | None / not reported |
| Angevaren et al. (2005) | Multiple | Older<br>(≥ 55 years) | Multiple | • Only cognitively healthy<br>• Measure of cardiorespiratory fitness | 11 | 9 | 82 | 0.55* | 0.17 [0.06, 0.29] | 4.86 | 78<br>(range 10–124) | 3.7<br>(range 2–6) | None / not reported |
| Barha et al. (2017) | Multiple | Older<br>(≥ 45 years) | Multiple | • Only cognitively and physically healthy<br>• Program duration ≥ 2 months<br>• More than one session per week<br>• Measure of memory, executive functions, verbal fluency, visuospatial ability, or processing speed | 39 | 32 | 302 | 0.71* | 0.19 [0.04, 0.34] | 73.91 | 125<br>(range 18–1476) | 7.4<br>(range 2–24) | None / not reported |
| Chen et al. (2020) | Multiple | Older<br>(≥ 55 years) | Executive functions | • Published between January 2003 and November 2019 | 32 | 16 | 94 | 0.21 | 0.16 [0.06, 0.25] | 28.13 | 57<br>(range 18–161) | 6.9<br>(range 1–24) | None / not reported |
| Colcombe & Kramer (2003) | Multiple | Older<br>(≥ 55 years) | Multiple | • Programs with an aerobic component | 18 | 8 | 82 | 0.31* | 0.17 [0.10, 0.25] | 6.03 | 85<br>(range 20–149) | 5.4<br>(range 2.5–12) | None / not reported |
| Falck et al. (2019) | Multiple | Older<br>(≥ 60 years) | Multiple | • Only cognitively and physically healthy<br>• Program duration > 2 months<br>• Measure of cardiorespiratory fitness | 58 | 24 | 223 | 0.24 | 0.15 [0.07, 0.22] | 32.28 | 105<br>(range 14–1476) | 6.1<br>(range 2–24) | Multilevel |
| Gasquoine & Chen (2020) | Multiple | Older | Executive functions | • Only cognitively and physically healthy<br>• Most used executive-functions neuropsychological tests: Digit Span backward, Digit Symbol, Trail-Making Test B, Letter fluency, and Color-Word Stroop<br>• Only non-physical exercise control activity | 26 | 21 | 106 | -0.09* | 0.12 [0.05, 0.19] | 26.39 | 118<br>(range 18–1476) | 7.4<br>(range 3–24) | Aggregates |
| Haverkamp et al. (2020) | Multiple | Adolescents & adults<br>(Mean 12–30 years) | Multiple | • Only cognitively healthy<br>• Controlled design, with or without random allocation | 28 | 13 | 65 | 0.36* | 0.39 [0.19, 0.60] | 72.20 | 116<br>(range 20–584) | 2.8<br>(range 1–6) | Aggregates |
| Hoffmann et al. (2021) | Aerobic | Older<br>(≥ 50 years) | Executive functions and episodic memory | • Only cognitively healthy<br>• Sedentary | 9 | 9 | 61 | 0.59* | 0.30 [0.06, 0.54] | 68.85 | 39<br>(range 24–120) | 5.9<br>(range 3–12) | Outcome selection |

| | | | | | | | | | | | | | |
|---|---|---|---|---|---|---|---|---|---|---|---|---|---|
| Jackson et al. (2016) | Multiple | Children & adolescents (7–12 years) | Executive functions | • Program duration ≥ 1 month | 8 | 2 | 7 | 0.20 | 0.06 [0.04, 0.07] | 60.66 | 136 (range 23–221) | 9 | Aggregates |
| Lindheimer et al. (2015) | Multiple | Adults & older | Multiple | • Studies with a physical control activity and a passive group<br>• Program duration ≥ 1 month | 9 | 1 | 22 | 0.37 | 0.09 [−0.42, 0.60] | - | 96 | 6 | Multilevel |
| Ludyga et al. (2020) | Multiple | All | Multiple | • Only cognitively and physically healthy<br>• Measure of complex attention, executive function or memory<br>• Program duration ≥ 1 month | 80 | 63 | 487 | 0.23 | 0.16 [0.12, 0.22] | 43.40 | 70 (range 15–584) | 5.7 (range 1–24) | Aggregates |
| Meli et al. (2021) | Multiple | Children (6–11 years) | Multiple | • Only cognitively and physically healthy<br>• Published between 2012 and 2021 | 6 | 2 | 6 | 0.65 | 0.12 [−0.13, 0.38] | 67.78 | 673 (range 158–931) | 2.4 (range 2–2.8) | Outcome selection |
| Northey et al. (2018) | Multiple | Older (≥ 50 years) | Multiple | • Without neurological or psychiatric conditions, except mild cognitive impairment<br>• Program duration ≥ 4 weeks | 43 | 29 | 191 | 0.29 | 0.18 [0.09, 0.26] | 39.65 | 52 (range 14–161) | 5.6 (range 1.5–12) | Multilevel |
| Rathore & Lom (2017) | Multiple | All | Working Memory | • Only cognitively and physically healthy | 15 | 5 | 12 | 0.27 | 0.25 [−0.09, 0.59] | 50.84 | 65 (range 36–100) | 3.7 (range 1–6) | Aggregates |
| Sanders et al. (2019) | Multiple | Older | Multiple | • Aged 18 years or older<br>• With and without cognitive impairment<br>• Program duration ≥ 1 month<br>• The physical intervention included a non-physical component<br>• Training intensity needs to be specified | 36 | 23 | 191 | 0.25* | 0.19 [0.10, 0.28] | 31.55 | 48 (range 16–119) | 5.5 (range 1–13) | Multilevel |
| Scherder et al. (2014) | Walking | Older (≥ 55 years) | Executive functions | • With and without cognitive impairment | 8 | 3 | 29 | 0.36 | 0.09 [−0.03, 0.20] | 55.90 | 115 (range 80–124) | 15 (range 6–24) | Aggregates |
| Smith et al. (2010) | Aerobic | Adults & older (≥ 18 years) | Multiple | • Only cognitively healthy<br>• Program duration > 1 month<br>• Not slow walking | 29 | 14 | 104 | 0.11* | 0.18 [0.10, 0.26] | 10.72 | 77 (range 10–149) | 5.5 (range 1.5–24) | Aggregates |
| Xiong et al. (2020) | Multiple | Older (Mean ≥ 60 years) | Executive functions | • Only cognitively healthy<br>• Program duration ≥ 1 month, ≥ 3 days/week, ≥ 20 min/session | 26 | 21 | 153 | 0.30* | 0.15 [0.07, 0.24] | 42.94 | 96 (range 15–1476) | 5.8 (range 1–24) | Outcome selection |
| Xue et al. (2019) | Multiple | Children & adolescents (6–17 years) | Executive functions | • Only cognitively and physically healthy<br>• Program duration ≥ 6 weeks | 19 | 9 | 26 | 0.23* | 0.30 [0.12, 0.48] | 96.91 | 320 (range 35–584) | 8 (range 1.5–24) | Outcome selection |
| Young et al. (2015) | Aerobic | Older (≥ 55 years) | Multiple | • Only cognitively healthy<br>• Measure of cardiorespiratory fitness<br>• Control activities: no treatment, a | 12 | 11 | 95 | 0.10* | 0.13 [0.02, 0.23] | 10.94 | 77 (range 10–124) | 5.5 (range 2–24) | None / not reported |

| | | | | | | | | | | | | |
|---|---|---|---|---|---|---|---|---|---|---|---|---|
| | | | | strength or balance program, or a program of social activities or mental activities | | | | | | | | |
| Zhao et al. (2022) | Multiple | Older (≥ 60 years) | Multiple | • Sedentary<br>• Only non-physical exercise control activity | 7 | 3 | 9 | 0.50 | 0.25 [−0.31, 0.81] | 63.40 | 36 (range 18–46) | 3.5 (range 3–4) | None / not reported |
| Zhidong et al. (2021) | Multiple | Older | Working memory | • Only cognitively healthy<br>• Session duration ≥ 10 min | 28 | 8 | 18 | 0.30 | 0.08 [−0.05, 0.21] | 0 | 69 (range 16–210) | 5.3 (range 1–12) | None / not reported |

* The overall effect size was the result of averaging the outcome of several meta-analyses reported in the article, each of them for specific cognitive domains.

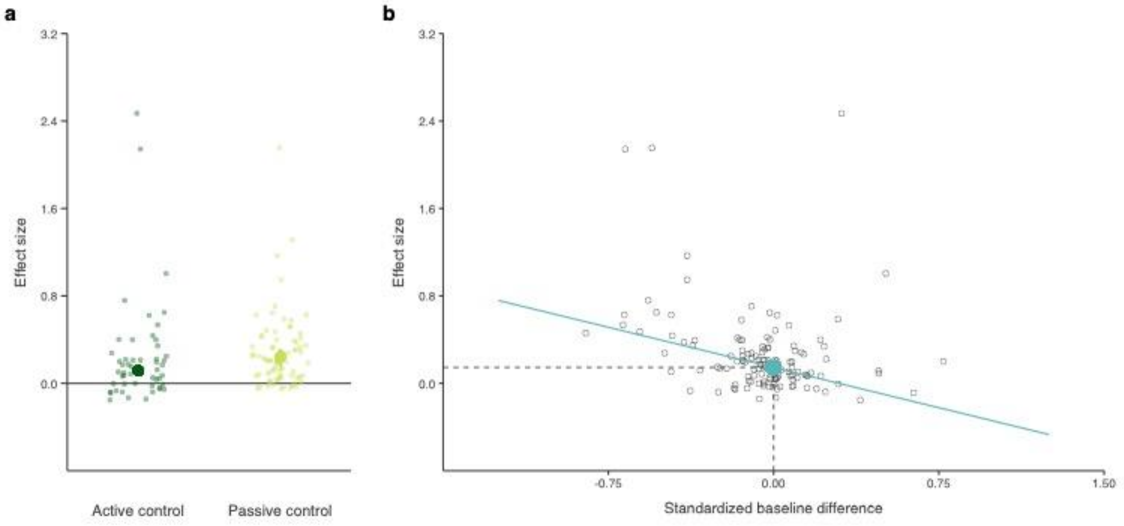


**Fig. S1**. Moderating influence of the type of control and baseline difference. Scatter plots of the observed effect sizes of primary studies as a function of (A) the type of control activity and (B) the between-group baseline difference. Passive controls and lower prior performance of the experimental group predicted larger effect sizes. Therefore, the studies with those characteristics were more likely to observe significant benefits than those with active controls and equivalent performance of both groups.
